# Baseline bioenergetic profile of mitochondria and permeabilized tissue of *Aedes aegypti* (Diptera: Culicidae) throughout its life cycle

**DOI:** 10.1101/2025.04.02.646709

**Authors:** Ruth Mariela Castillo-Morales, Diana Lizeth Urbina-Duitama, Stelia Carolina Mendez-Sanchez, Jonny E Duque

## Abstract

Cellular respiration parameters provide information on the energy metabolism of an organism under physiological or non-conditions and can be determined by isolating mitochondria or even using their permeabilized tissue. The present study aimed to describe the mitochondrial activity of *Ae. aegypti* throughout its life cycle. For this purpose, we isolated mitochondria and permeabilized tissue (1 and 5 individuals) from larvae (L4), pupae, and adults (female thorax) to estimate oxygen consumption using different oxidizable substrates (glutamate, malate, proline + pyruvate, succinate, and glycerol-3-phosphate). The Respiratory Control Coefficient (RCC) was calculated, and the specific enzyme activity of the Electron Transport Chain (ETC) was quantified. The highest values of oxygen consumption (state 3) were obtained when succinate was used as the oxidizable substrate in all developmental stages of *Ae. aegypti*, with a progressive increase in the values recorded from larva (isolated mitochondria: 30.3 ± 5.0 pmol/(s*mL), permeabilized tissue/ 5 individuals: 48.8±1.8 pmol/(s*mL)), through pupa (isolated mitochondria: 42.8±3.6 pmol/(s*mL); permeabilized tissue/5 individuals: 47.4±2.2 pmol/(s*mL)), through adult (isolated mitochondria: 53.4±6.0 pmol/(s*mL); permeabilized tissue/5 thorax: 27.4±2.4 pmol/(s*mL)). These results are congruent with the values obtained for enzyme activity, with higher activity of succinate oxidase and succinate dehydrogenase enzymes (Complex II). The CCR values for isolated mitochondria and permeabilized tissue ranged from 1 to 1.8. The results allow us to conclude that there are changes in mitochondrial respiration throughout the life cycle of the *Ae. aegypti* mosquito.

## Introduction

The mitochondria are an essential subcellular organelle for eukaryotic cells by synthesizing ATP through oxidative phosphorylation, actively participating in cell signaling, and synthesizing key molecules in cell functioning (Picard et al., 2018). In terms of morphological parameters, the size and shape of mitochondria are variable because they can divide and fuse according to different physiological stages of development, energy requirements of the cell, and in response to external stimuli (Giacomello et al., 2020; Palmer et al., 2011; Popkov et al., 2016).

One of the main functions of mitochondria lies in oxidative phosphorylation. This process occurs in the mitochondrial respiratory chain, and measuring it provides data into the physiology of this organelle about the different states of its molecular components, enzyme levels, metabolites (Nicholls et al., 2012; Giacomello et al., 2020). Additionally, it helps in understanding changes associated with metabolic alterations and the activity of the enzymes in the electron transport chain (Giacomello et al., 2020).

In this context, the study of mitochondria in insects has primarily focused on understanding their physiology and quantifying respiratory parameters to be used as a target point of action of new molecules (whether synthetic or of natural origin) and description of potential mechanisms of resistance to insecticides (Bajda et al., 2017; Bolter & Chefurka, 1990; Colinet et al., 2017; Giulivi et al., 2008; Hu et al., 2008).

For *Ae. aegypti*, the primary vector of arboviruses diseases like dengue, Zika and chikungunya (Rodriguez, 2002; Bhatt et al., 2013; Castrillón et al., 2014; Rodriguez-Morales, 2015a, 2015b; Weaver and Lecuit, 2015), publications related to mitochondrial respiratory measurement initially focused on the description of isolation protocols to optimize the measurement of different respiratory parameters under normal and stress conditions (Picard, Taivassalo, Gouspillou, et al., 2011; Picard, Taivassalo, Ritchie, et al., 2011). Another part of the research describes the effects of blood feeding on functional (mitochondrial respiration, electron transport chain activity) and structural parameters of the female thorax, where oxygen consumption values with blood feeding (proline elicitation) are slightly lower than those obtained with sugar-based feeding (glucose elicitation) (Correa et al., 2015; Gaviraghi & Oliveira, 2019; Goncalves et al., 2009). In recent years, some studies have quantified mitochondrial physiology and the functioning of protein complexes at the electron transport chain in response to compounds with insecticidal activity, such as essential oils, plant extracts, or their metabolites (Borrero et al., 2018; Borrero-Landazabal et al., 2020; Castillo-Morales et al., 2019).

Given that the assessment of mitochondrial function is crucial for understanding how different biological conditions or xenobiotics affect its activity (Palmeira & Rolo 2011), the present study aim is to describe, for the first time, the mitochondrial respiratory behavior throughout the life cycle of *Ae. aegypti*. The description of these respiratory parameters can contributes to the understanding of the physiological and biochemical mechanisms of *Ae. aegypti* throughout its life cycle. These data could be used to improve the evaluation of the mechanisms of action of new molecules for the control of insects of public health interest.

## Methodology

### Biological material

Individuals from a colony of *Aedes aegypti* (Rockefeller strain) were maintained under insectary conditions at 25 ± 5 °C, with a relative humidity 70 ± 5 %, and a photoperiod 12:12 hours. Adults mosquitoes were permanently fed with a sugar solution made from approximately 10% honey. To obtain eggs from the female mosquitoes, they were fed with blood from albino rats (*Rattus norvegicus*) of the Wistar WI IOPS AF/Han breed, provided by the biotherium of the Universidad Industrial de Santander. All experimental processes were under prior approval of the Ethics Committee under minute No. 21 of November 23, 2018. The oviposits obtained were submerged in dechlorinated water with some TetraMin Tropical Flakes® fish concentrate to stimulate hatching. Hatched larvae were kept in plastic trays and fed with the same concentrate that stimulated hatching.

### Mitochondria isolation

Mitochondria were isolated following a modified version of the protocol described by Borrero et al., (2018). Specimens were selected from the colony according to treatment, including approximately 2000 L3-L4 larvae and 300 pupae, and 180 adult females (Flight muscle) without prior blood feeding, using the protocol of Gaviraghi and Oliveira (2019).

Once the individuals were separated into the developmental stages of interest for the study, each group was homogenized in a Potter-Elvehjem with 10 mL of isolation medium at 4◦C, composed of sucrose 250 mM, HEPES 10 mM, EGTA 1 mM, and bovine serum albumin (BSA) at 0.1% w/v, pH 7.4. The preparation was maintained at 4°C through subsequent washes and centrifugations. The product obtained from homogenization was filtered through glass wool to remove chitin and tissue debris that did not go through an adequate maceration process. The resulting solution was subjected to four centrifugations, the first at 300 xg for 5 minutes and the sediment was discarded. The second centrifugation was performed at 8000 xg for 10 minutes, where the supernatant was discarded, and the sediment was gently resuspended in 10 mL of the same isolation medium. The third centrifugation was performed at 7000 xg for 10 minutes, discarding the supernatant, and the sediment was resuspended in 10 mL of isolation medium without BSA. The fourth centrifugation was performed at 7000 xg for 10 minutes, discarding the supernatant. The final pellet was suspended in 250 μL of isolation medium without BSA and stored at 4 °C. Bradford’s colorimetric method was used to determine the total protein concentration of the isolated mitochondria (Bradford, 1976). The absorbance measurement was performed with a Multiskan Go spectrophotometer (Thermo Scientific) at a wavelength of 595 nm (Bradford, 1976).

### Measurement of respiration in isolated mitochondria

Mitochondrial respiration at each developmental stage of *Ae. aegypti* was measured polarographically using a high-resolution, two-channel oxygraph (Oxygraph-2k, Oroboros Instruments, Innsbruck, Austria) at 27.5 °C and 750 rpm. Each sample was placed in a 2 mL chamber containing an incubation medium composed of 100 mM HEPES, 0.1 mM EGTA, 125 mM D-Mannitol, and 65mM KCl (pH 7.4). Forr all developmental stages (larvae between L3-L4, pupae and thoraces), 0.05 mg/mL of total mitochondrial protein was added. Oxygen consumption was evaluated in response to oxidizable substrates including 5 mM sodium glutamate, 10 mM malate, 10 mM proline + 10 mM sodium pyruvate, 3 mM sodium succinate, and 20 mM glycerol-3-phosphate. For the succinate and glycerol-3-phosphate substrates, 1 μM rotenone was previously added to inhibit the passage of electrons through complex I of CTE and ensure their entry through complex II and glycerol-3-phosphate dehydrogenase.

KH2PO4 (1.6 mM) was included, and the oxygen consumption in this basal condition rate was recorded as state 2. The highest respiration rate (state 3) was achieved following the adding 0.6 mM ADP to. With the addition of Oligomycin 3.4 μM (inhibitor of the F0 subunit of ATP synthase), a respiratory state of low O2 consumption was induced, called induced state 4. Finally, 1 μM FCCP, an uncoupling agent, was added to determine the uncoupled respiration rate. Oxygen consumption rates were expressed as pmol*sec-1*106 O2 consumed x mg-1 protein, and were recorded in real time using DatLab 4.0 software (Oroboros Inc., Austria).

### Respiration measurement on larvae, pupae and permeabilized females thoraces

Oxygen consumption was measured in permeabilized tissue of each developmental stage of *Ae. aegypti*. A modified version of the protocol by Gaviraghi and Oliveira (2019) was employed. One or five individuals were placed separately in each each chamber of the oxygraph, containing 2 mL of respiration medium. For thoracic measurement in females, the sclerites of each were carefully opened with fine-tipped forceps to expose the flight muscle prior to their addition to the chamber. Respiration was measured under previously described conditions,, with the exception of the inclusion of 0.05 mg/mL digitonin, a non-ionic detergent used to permeabilize cell membranes, facilitating the passage of substrates that promote oxidative phosphorylation. Measurements were initiated following signal stabilization in the oxygraph (Correa et al., 2012; Vercesi et al., 1991).

### Calculation of the Respiratory Control Coefficient (RCC)

The Respiratory Control Coefficient (RCC) is a parameter for testing the integrity of a mitochondrial preparation. It is defined as the value obtained by dividing the rate of oxygen consumption observed when there is maximal ATP synthesis (i.e. in the presence of ADP or the presence of a proton translocator), and by the rate of oxygen consumption when there is no ATP synthesis or in the absence of a proton translocator.

The RCC was calculated for all developmental stages (L4 larvae, pupae, and adult female thoraces) and for each substrate evaluated, using the formula: Respiration stage 3 / Respiration stage 4 induced with oligomycin. Regarding the action of oligomycin, it is a reagent that binds to the F0 proton channel of ATP synthase, blocking proton translocation, and ATP synthesis and inhibiting O2 uptake (Hearne et al. 2020). Since oligomycin uncouples the electron transport chain by blocking ATP synthesis, measuring oxygen consumption under these conditions provides valuable insights into mitochondrial function, bioenergetic capacity, and efficiency (Correa et al., 2015).

### Measurement of the specific enzyme activity of CTE

To understand the functioning of the mitochondrial respiratory chain at different developmental stages of *Ae. aegypti*, the activity of eight enzymes associated with the four respiratory complexes and ATPase was evaluated. For this purpose, mitochondria isolated from larvae (L4), pupae, and thorax of adult female *Ae. aegypti* and fragmented through sonication cycles of 15 minutes at 4°C. Enzyme activity was measured by respirometry using a high-resolution oxygraph (Oxygraph-2k, Oroboros Instruments, Innsbruck, Austria) or by spectrophotometry (UV/Vis Multiskan GO spectrophotometer from Thermo Scientific). The activities of NADH oxidase, succinate oxidase,

NADH dehydrogenase, and succinate dehydrogenase were quantified using the method described by Singer (Singer, 1974), while NADH cytochrome c reductase and succinate cytochrome c reductase activities were measured by the method described by Somlo (Somlo, 1965). The activity of NADH cytochrome c reductase and cytochrome c oxidase followed the methodology described by Mason & Schatz (Mason & Schatz, 1973). Finally, ATPase activity in fragmented mitochondria was assessed using the methodology described by (Pullman et al. 1960).

### Statistical analyses

All the experiments were conducted in triplicate across independent trials, with each concentration evaluated three or four times. Data were tabulated and analyzed using descriptive statistics and Kolmogorov-Smirnorv and Shapiro Wilk normality tests. For data following a normal distribution, ANOVA followed Tukèy tests was applied. When the data did not meet normality assumtions, non-parametric tests were employed (Kruskal-Wallis), and if they were significant (p≤0.05), multiple comparison tests were used. The results were analyzed with Statistics V11 and GrahPad Prisma 8.0.

## Results

### Oxygen consumption in mitochondria isolated from L4 larvae

The highest rates of oxygen consumption in isolated mitochondria from L3-L4 larvae were observed following addition of the succinate substrate. The oxygen consumption values for stage 3 were 30. 3±5 pmol/(s*mL), state 4 was 28.4±5 pmol/(s*mL), induced state 4 was 22±7.6 pmol/(s*mL) and in the presence of FCCP, 13.62±6.3 pmol/(s*mL) (Figure 1). Significant statistical differences were found when comparing oxygen consumtion between substrates in each respiratory state. Differences were noted between succinate and other substrates in state 3 (KW test H: (4, N=72) =54.766580 p=0.003167), state 4 (KW test H: (4, N=72) =56.11636 p=0.005921), state 4 induced (KW test H: (4, N=72) =44.54037 p=0.00000) and FCCP respiration (KW test H: (4, N=56) =16.15322 p=0.003789). Although complex II does not pump protons across the mitochondrial membrane, a higher rate of oxygen consumption was observed with succinate compared to the other substrates. This elevated activity may reflect increased energy requirement and biomass acquisition in response to varying environmental conditions in larvae. The oxygen consumption in larvae with succinate suggest higher complex II activity, likely linked to the metabolic demands for biomass acquisition and energy storage during larvae and pupae stages.

**Figure 1.**
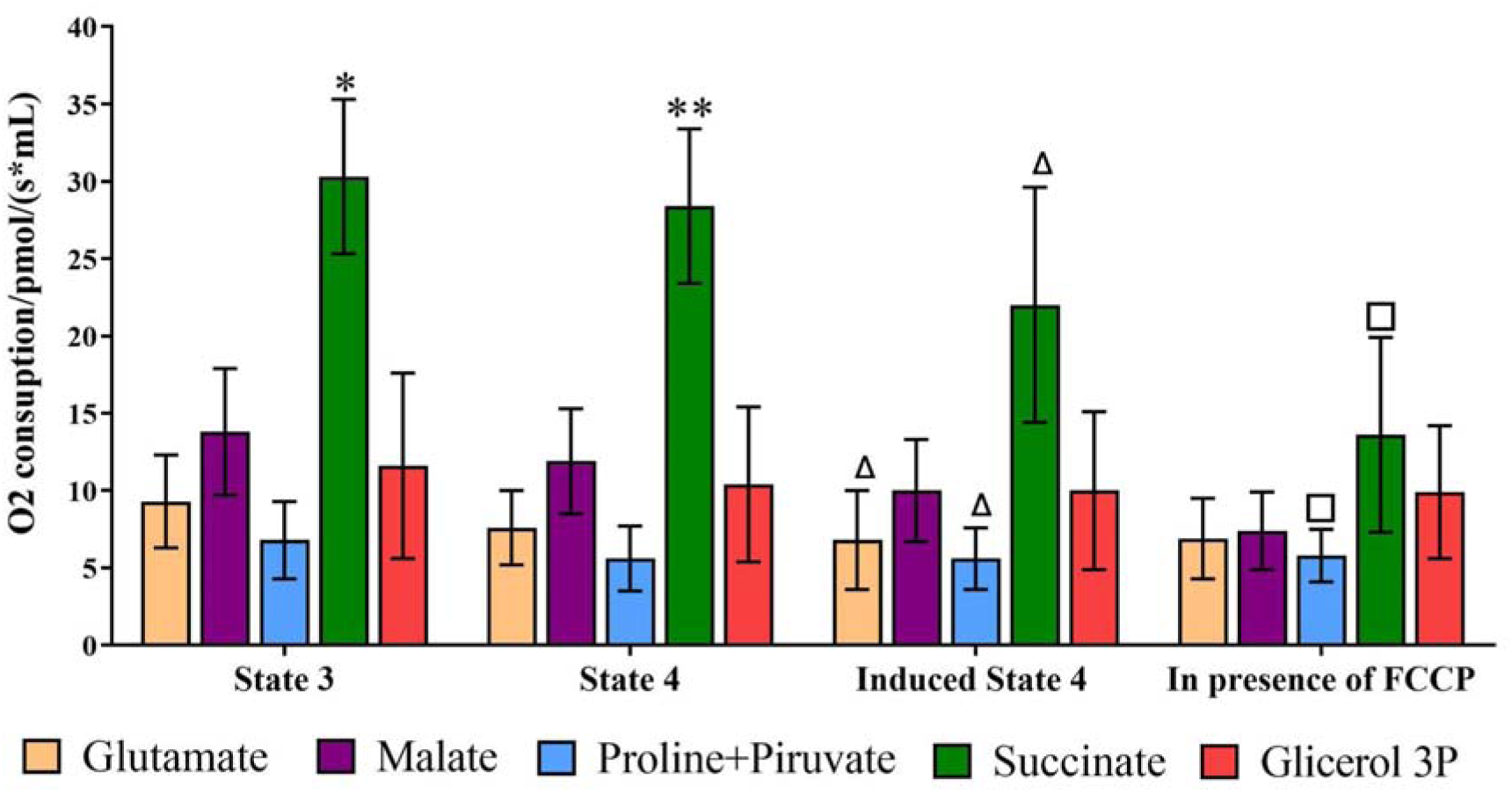
Average oxygen consumption (pmol/(sec*mL)) of mitochondria isolated from L3-L4 larvae of *Ae. aegypti* for respiratory stages: stage 3, stage 4, oligomycin-induced stage 4, and respiration in the presence of FCCP (uncoupled mitochondria). The oxidizable substrates used were glutamate, malate, proline+pyruvate, succinate, and G3P. Different symbols indicate significant statistical differences between *Oxidizable substrates for state 3. **Oxidizable substrates for state 4. Δ Oxidizable substrates for induced state 4. □ Oxidizable substrates in the presence of FCCP (KW test 2 p≤0.05).

### Oxygen consumption of pupal mitochondria

Similar to larval mitochondria, increased oxygen consumption was observed in pupae when succinate was used as the substrate across all respiratory stages: stage 3 of 42.8±3.6 pmol/(s*mL), stage 4 of 39.5±4.5 pmol/(s*mL), induced stage 4 of 36.4±3.7 pmol/(s*mL) and respiration in the presence of FCCP of 28.5±8.0 pmol/(s*mL). Significant differences in oxygen consumption were detected when comparing the respiratory states using other substrates, such as glutamate, malate, proline+pyruvate, and G3P. These differences were statistica significant in state 3 (KW test H: (4, N=39) =32.76122 p=0.00255) and state 4 (KW test H: (4, N=39) =31.85306 p=0.000106). The same trend was observed in induced state 4 (KW test H: (4, N=39) =27.6700 p=0.000184) and respiration in the presence of FCCP (KW test H: (4, N=39) =24.77551 p=0.000253) (Figure 2).

**Figure 2.**
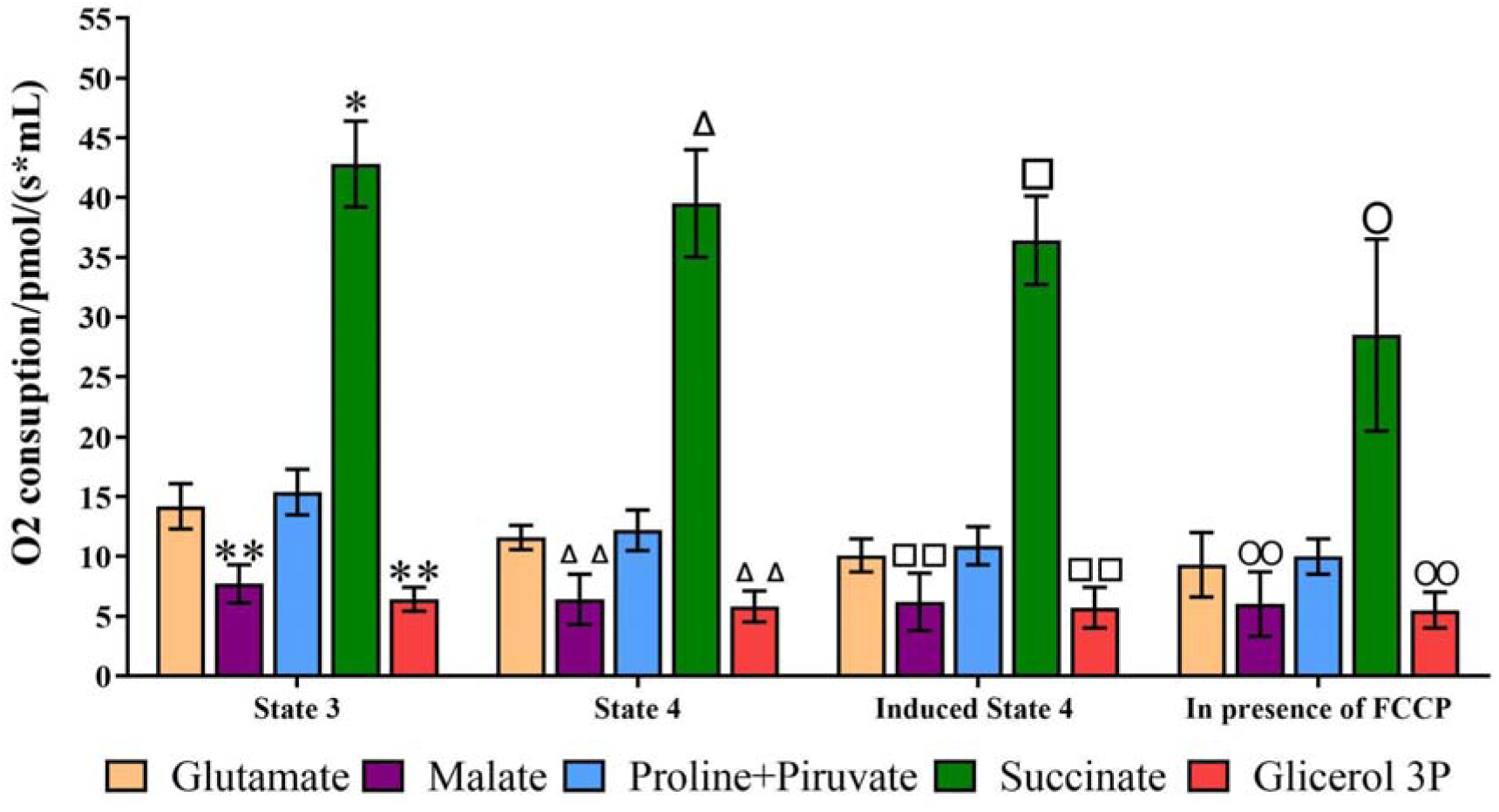
Average oxygen consumption (pmol/(sec*mL)) of mitochondria isolated from *Aedes aegypti* pupae across various respiratory states: state 3, state 4, state 4 Induced with Oligomycin and respiration in the presence of FCCP (uncoupled mitochondria). The oxidizable substrates utilized include glutamate, malate, proline+pyruvate, succinate, and G3P. Different symbols indicate significant statistical differences between *, ** Oxidizable substrates for state 3; Δ, ΔΔ Oxidizable substrates for state 4; □, □□ Oxidizable substrates for state 4 induced; □, □□ Oxidizable substrates for state in the presence of FCCP (KW test p≤0.05).

### Oxygen consumption of mitochondria isolated from adult female thorax

Mitochondria isolated from the thorax adult female exhibited higher oxygen consumption across all respiratory states when using succinate as the oxidizable substrate. The measured oxygen consumption values were as follow: state 3 of 53.4 ± 6.0 pmol/(s*mL), state 4 of 51.5 ± 7.8 pmol/(s*mL), induced state 4 of 51.1 ± 6.0 pmol/(s*mL) and respiration in the presence of FCCP of 53.5±1.8 pmol/(s*mL). Comparative analysis revealed significant differences in oxygen consumption betwen succinate and others substrates evaluated. Specifically, significant differences were observed for stage 3 between glutamate, malate, proline+pyruvate and G3P. Stage 3 (KW test H: (4, N=44) =37.65303 p=0.000287), state 4 (KW test H: (4, N=44) =36.27515 p=0.046388), induced state 4 (KW test H: (4, N=44) =35.52607 p=0.019768) and respiration in the presence of FCCP (KW test H: (4, N=38) =25.89530 p=0.016646).

The second substrate with the highest oxygen consumption in female thorax was G3P. Statistical differences were observed when comparing G3P to the other substrates evaluated. Specifically, significant differences were found between G3P-glutamate and G3P-malate at stage 3(KW test H: (4, N=44) =37.65303 p≤0.05). Similar differences were noted at stage 4 (KW test H: (4, N=44) =36.27515 p≤0.05) and induced state 4 (KW test H: (4, N=44) = 35.52607 p≤0.05) Aditionally, significant were identifitied among G3P-glutamate, G3P-malate and G3P-proline+pyruvate for respiration in the presence of FCCP (KW test H: (4, N=38) =25.89530 p≤0.05) (Figure 3).

**Figure 3.**
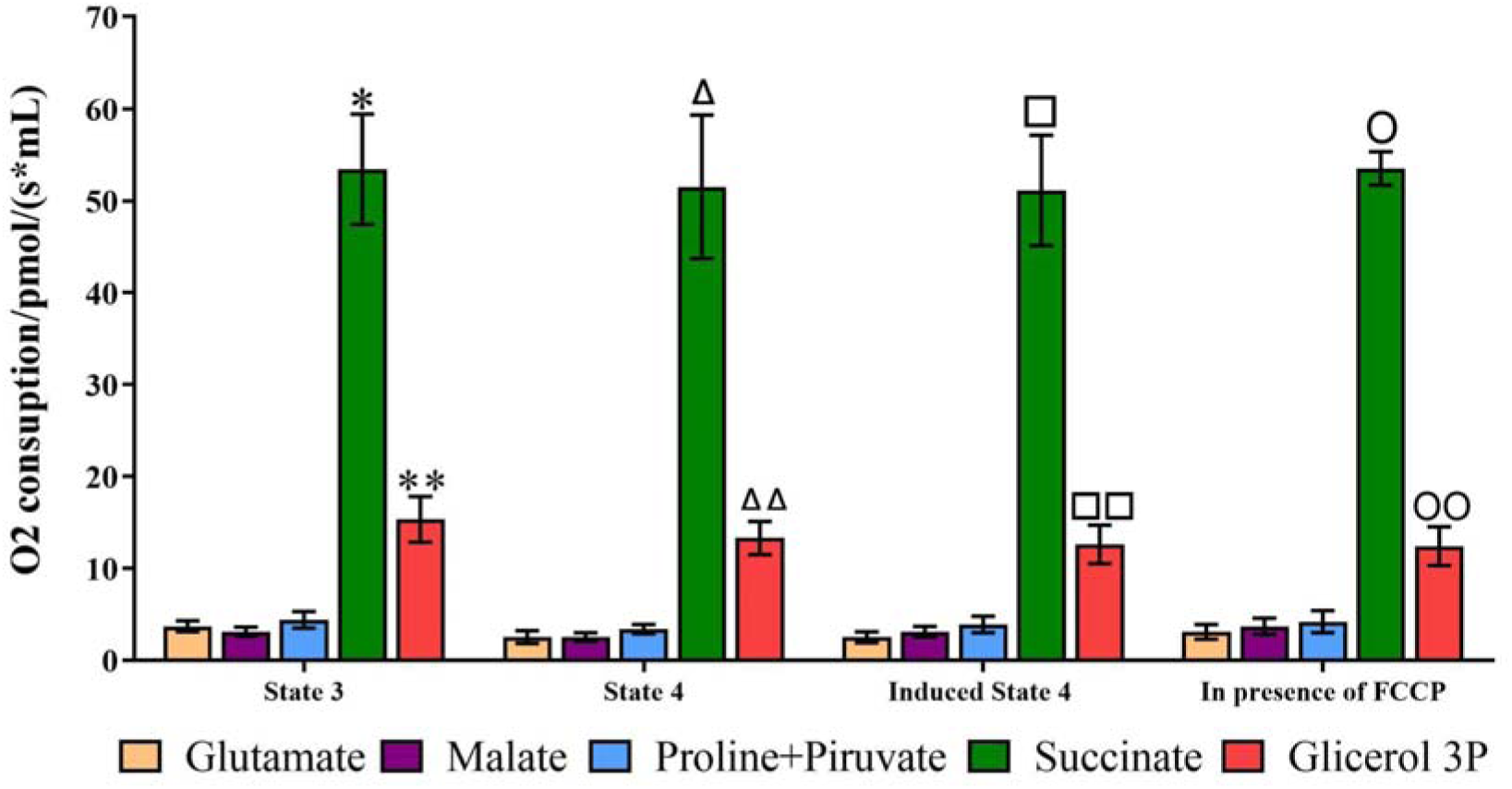
Average oxygen consumption (pmol/(sec*mL)) of mitochondria isolated from the thorax of *Ae. aegypti* females for respiratory states: state 3, state 4, and oligomycin - induced state 4, as well as respiration in the presence of FCCP (uncoupled mitochondria). The oxidizable substrates glutamate, malate, proline+pyruvate, succinate, and G3P were used. Different symbols indicate significant statistical differences between *, **Oxidizable substrates for state 3; Δ, ΔΔ Oxidizable substrates for state 4; □, □□ Oxidizable substrates for induced state 4; 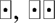 Oxidizable substrates in the presence of FCCP (KW test p≤0.05).

### Comparison of oxygen consumption of isolated mitochondria between developmental stages

When comparing the respiratory values at stage 3 for each oxidizable substrate and each developmental stage, it was observed that the highest values were significantly recorded with the succinate substrate (Supplementary Material - Figure 1). As the life cycle of the mosquito changes stage, a progressive increase in oxygen consumption was detected with this substrate, starting with larva (30.3 ± 5.0 pmol/(s*mL)) and reaching a maximum value in adult (53.4 ± 6.0 pmol/(s*mL)). When comparing oxygen consumption values between developmental stages, significant differences were found between L4 larva vs pupa, L4 larva vs adult, and pupa vs adult (ANOVA test F=61.629, df=2, p≤0.05) (Supplementary Material - Figure 1).

With the other substrates, much lower oxygen consumption was observed, with similar behavior for glutamate and proline+pyruvate substrates, with higher respiratory values in pupae (glutamate: 14.2±1.9 pmol/(s*mL); proline+pyruvate: 15.4 ± 1.9 pmol/(s*mL)) compared to larval stages (glutamate: 9.3 ± 3.0 pmol/(s*mL); proline+pyruvate: 6.8 ± 2.5 pmol/(s*mL)) and adult (glutamate: 3.7 ± 0.6 pmol/(s*mL); proline+pyruvate: 4.4 ± 0.9 pmol/(s*mL)), with significant differences for glutamate substrate between L4 larva vs pupa (ANOVA test F=49.1980, df=2, p= 0.000199); L4 larva vs adult (ANOVA test F=49.1980, df=2, p= 0.00012); pupa vs adult (ANOVA test F=49.1980, df=2, p= 0.000121). With the substrate proline+pyruvate, significant statistical differences were found between larva L3-L4 vs pupa, larva L3-L4 vs adult, and pupa vs adult (ANOVA test F=58.5959, df=2, p≤0.05) (Supplementary Material - Figure 1).

### Respiratory Control Coefficient (RCC)

In the present study, the CCR was calculated for all developmental stages (L3-L4 larvae, pupae and female thorax) and for each substrate evaluated as follows: Respiration stage 3 / Respiration stage 4 induced with oligomycin. The oligomycin-induced CCR values obtained for all mitochondrial isolates showed values ranging from 1 to 1.6 (Table 1). For larval L4 mitochondria, the highest values of induced CCR were observed with the substrates glutamate (1.6 ± 0.7) and succinate (1.6 ± 0.9). When comparing the values between the substrates evaluated, no significant statistical differences were found between them (KW test H: (4, N=72) = 5.167793 p=0.2705). For pupal mitochondria, the highest values of induced CCR were found for the substrates glutamate (1.4 ± 0.3) and proline+pyruvate (1.4 ± 0.1), with significant statistical differences between them (KW test H: (4, N=39) = 10.21238 p=0.0370). Finally, for thorax mitochondria, the highest value was obtained with the glutamate substrate (1.6 ± 0.5), and significant statistical differences were found between glutamate and malate substrates (KW test H: (4, N=44) = 16.43407 p=0.0001325) (Table 1).

**Table 1.** Respiratory Control Coefficient (RCC: Respiration state 3 / Respiration state 4 induced with oligomycin) in mitochondria isolated from L4 larvae, pupae, and thorax of *Ae. aegypti* females. Mitochondria were assessed using oxidizable substrates: glutamate, malate, proline+pyruvate, succinate, and glycerol-3 phosphate (G3P). Significant statistical differences among substrates are indicated by different letters: a,b denote differences between glutamate and proline+pyruvate in mitochondria isolated from pupae ((KW test H: (4, N=39) = 10.21238 p=0.0370), c,d show differences between glutamate and malate for mitochondria isolated from the thorax (KW test H: (4, N=44) = 16.43407 p=0.0001).

| Stage of development | Induced CCR value ± Deviation | | | | |
| --- | --- | --- | --- | --- | --- |
|  | Glutamate | Malate | Proline+Piruvate | Succinate | G3P |
| **Larvae** | 1,6 ± 0,9 | 1,4 ± 0,3 | 1,3 ± 0,3 | 1,6 ± 0,7 | 1,2 ± 0,1 |
| **Pupae** | 1,4 ± 0,3 <sup>a</sup> | 1,3 ± 0,3 | 1,4 ± 0,1 <sup>b</sup> | 1,2 ± 0,1 | 1,2 ± 0,2 |
| **Thorax** | 1,6 ± 0,5 <sup>c</sup> | 1,0 ± 0,2 <sup>d</sup> | 1,2 ± 0,1 | 1,1 ± 0,2 | 1,2 ± 0,2 |

### Measurement of respiration in larvae, pupae, and thorax of permeabilized females

The average respiratory values at stage 3 were compared for each oxidizable substrate and each developmental stage, showing differences between the values obtained for 1 and 5 permeabilized individuals. With a single individual, the highest oxygen consumption for larvae were recorded with the substrate glutamate (8.8 ± 1.9 pmol/(s*mL)) and succinate (8.0 ± 1.7 pmol/(s*mL)); for pupae it was the substrate proline+pyruvate (9.7 ± 0.9 pmol/(s*mL)) and succinate (9.3 ± 0.8 pmol/(s*mL)); and for adult thorax it was the substrate succinate (12.6 ± 1.1 pmol/(s*mL)).

Additionally,No consistent trend in oxygen consumption was observed across developmental stages. Upon statistical comparison between stages for each substrate used, significant differences were found for the substrate glutamate between Larva L4 vs adult and pupa vs adult (KW test H: (2, N=29) =19.11232 p≤0.05); for malate between Larva L4 vs adult and pupa vs adult (Tukey test F=56.022, df=2, p≤0.05); for proline+pyruvate between Larva L4 vs adult and pupa vs adult (KW test H: (2, N=28) =22.91690 p=≤0.05); for succinate between Larva L4 vs adult and between pupa vs adult (ANOVA: F=27.266, df=2, p≤0.05); and for G3P between Larva L4 vs pupa and between pupa vs adult (ANOVA: F=8.2923, df=2, p≤0.05) (Figure 4a).

**Figure 4.**
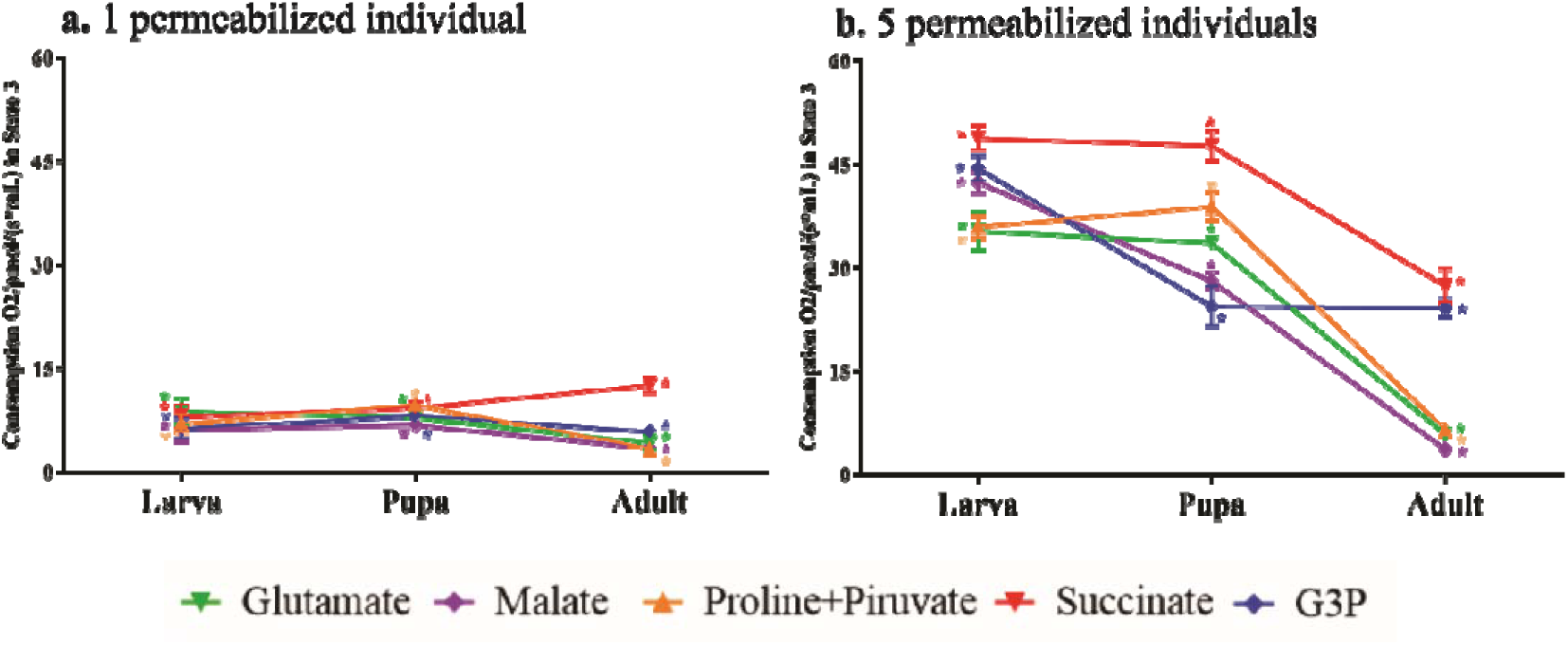
Comparison of average oxygen consumption (pmol/(sec*mL)) at stage 3 of permeabilized individuals of larvae, pupae, and thorax of *Ae. aegypti* females for the oxidizable substrates glutamate, malate, proline+pyruvate, succinate and G3P. a. Permeabilized individuals. *Significant statistical differences between larvae, pupae and adults for each oxidizable substrate evaluated: glutamate (KW test p≤ 0.005); malate (ANOVA test p≤ 0.005); proline+pyruvate (KW test p≤0.05); succinate (ANOVA test p≤0.05) and G3P (ANOVA test p≤0.05). b. Five permeabilized individuals. *Statistically significant differences between larvae, pupae and adults for each oxidizable substrate evaluated: glutamate (KW test p≤ 0.005); malate (KW test p≤ 0.005); proline+pyruvate (KW test p≤0.05); succinate (ANOVA test p≤0.05) and G3P (KW test p≤0.05).

On the contrary, in groups of the five individuals, the highest oxygen consumption across all stages evaluated was presented with the succinate substrate, remaining almost stable in larva (48.8 ± 1.8 pmol/(s*mL)) and pupa (47.4 ± 2.2 pmol/(s*mL)), but decreasing drastically in adult (27.4 ± 2.4 ±2.4 pmol/(s*mL)). For this substrate, significant differences were found between Larva L4 vs adult and between pupa vs adult (KW test H: (2, N=19) =12.32556 p≤0.05) (Figure 4b). Oxygen consumption with the substrates glutamate and proline+pyruvate presented a similar trend in the two substrates, with a slight increase from larva to pupa, and then a drastic decrease in the adult stage. For the substrate glutamate, significant differences were found between Larva L4 vs adult and between pupa vs adult (KW test H: (2, N=22) =14.76920 p≤0.05); for the substrate proline+pyruvate between Larva L4 vs adult (KW test H: (2, N=23) =19.47826 p=0.000034) (Figure 4b).

With the malate substrate, high oxygen consumption was observed in larva stage (42.4 ± 1.6 pmol/(s*mL)), which decreased sharply in the pupal (28.1 ± 1.3 pmol/(s*mL)) and adult stage (3.7 ± 0.5 pmol/(s*mL)), with significant differences between Larva L4 vs adult and between pupa vs adult (KW test H: (2, N=23) =19.21739 p≤0.05). For the G3P substrate, high values of oxygen consumption were presented in larvae (4.4 ± 1.9 pmol/(s*mL)), followed by a marked decrease in pupa (24.4 ± 2.9 pmol/(s*mL)), with levels remaining stable in adult (24.2 ± 1.4 pmol/(s*mL)), with significant differences between Larva L4 vs pupa and Larva L4 vs adult (KW test H: (2, N=19) =13.24632 p≤0.05) (Figure 4b).

The use of succinate substrate was predominant in all developmental stages, suggesting a greater input of electrons via Complex II. However, in juvenile stages (larvae and pupae), notable oxidation of substrates such as glutamate, malate, and proline+pyruvate was notorious, indicating three electron entry pathways: via Complex I and II for larvae, and via Complex II and dehydrogenases (Ubiquinone) for pupae. In adults, a marked preference for electron entry via Complex II and dehydrogenases (Ubiquinone) was observed, in comparison with the values observed with the other substrates evaluated. It is worth noting that the results obtained with five permeabilized individuals were like the results obtained with isolated mitochondria. Additionally, the differences between the values obtained for each substrate are observable with five individuals, facilitating the comparison and analysis of the results. Considering this fact, the results suggest that the evaluation of respiratory values should preferably be performed with 5 permeabilized individuals.

### Calculation of Respiratory Control Coefficient (RCC) induced in permeabilized tissue

In permeabilized larvae, the highest induced RCC were obtained with a single permeabilized larva, particularly with the substrates glutamate (1.5 ± 0.4) and succinate (1.5 ± 0.3), with significant differences between glutamate and G3P 4 (ANOVA: F= 3.043, df=4, p=0.031737), and between succinate and G3P (ANOVA: F= 3.043, df=4, p=0.040495). For five permeabilized larvae, the highest CCR value was observed with the glutamate substrate (1.3 ± 0.1), with significant differences between glutamate-succinate, malate-succinate, and succinate-G3P substrates (KW test H: (4, N=36) = 20.24180 p≤0.05).

For permeabilized pupae, the highest values of induced CCR were obtained for one pupa permeabilized with the G3P substrate (1.6 ± 0.2), with significant differences between the substrates glutamate-succinate, glutamate-G3P, malate-G3P, proline+pyruvate-G3P and succinate-G3P (ANOVA: F= 10.381, df=4, p≤0.05). For five permeabilized pupae, the highest value of induced CCR was found for the substrate proline+pyruvate (1.4 ± 0.1), finding significant differences between the substrates glutamate-succinate, malate-succinate, proline+pyruvate-succinate (ANOVA: F= 10.381, df=4, p≤0.05) (Table 2).

**Table 2.**
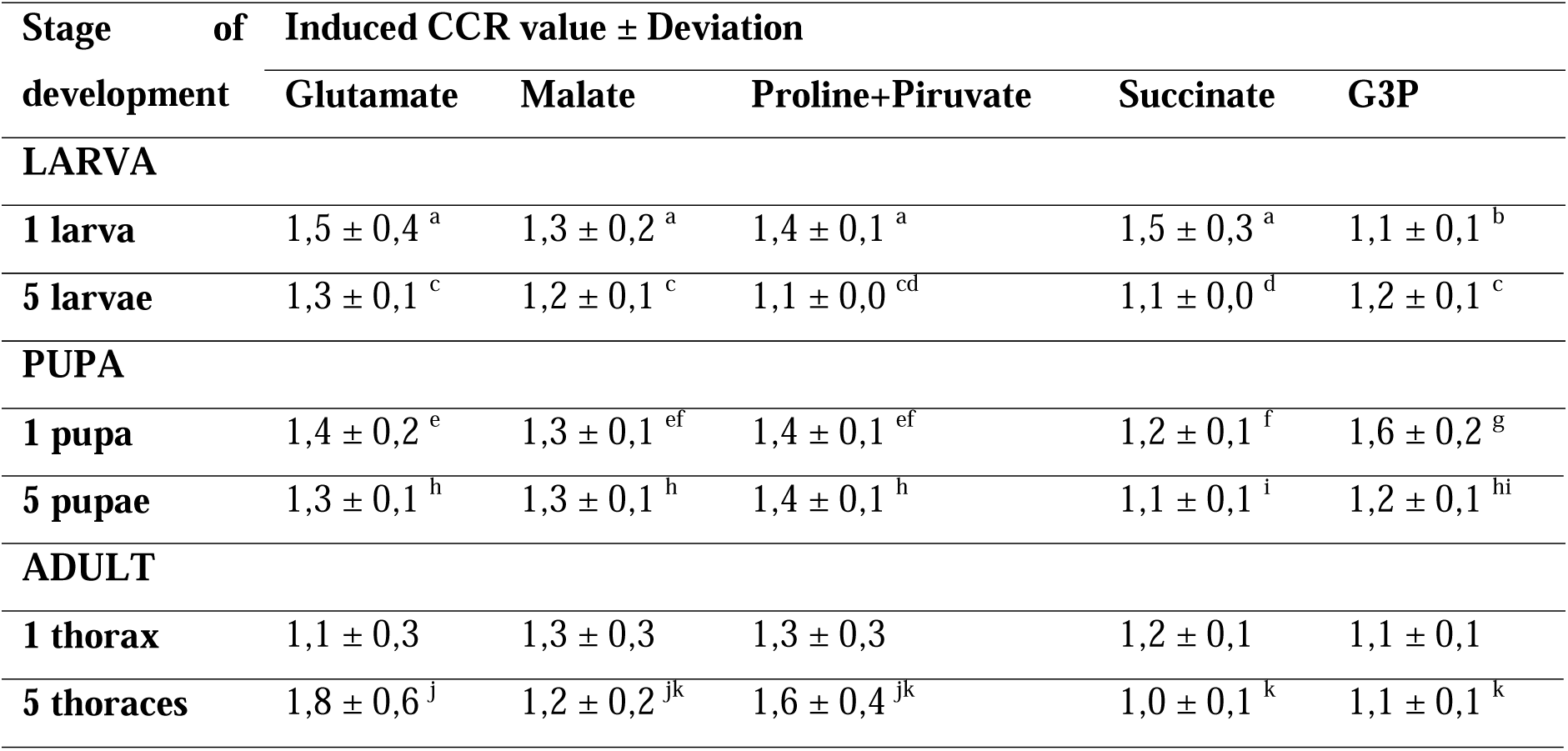
Respiratory Control Coefficient (RCC: Respiration state 3 / Respiration state 4 induced with Oligomycin) of permeabilized individuals of L4 larvae, pupae and thorax of *Ae. aegypti* females. The oxidizable substrates glutamate, malate, proline+pyruvate, succinate and G3P were used. Different letters indicate significant statistical differences for: a,b one larva between substrates (ANOVA test p≤0.05); five larvae between substrates a,b (KW test p≤0.05; c,d five larvae between substrates (KW test p≤0.05); e,f,g one pupa between substrates (ANOVA test p≤0,05); h,i five pupae between substrates (ANOVA test p≤0,05); j,k five thoraxes between substrates (KW test p≤0,05).

| Stage of development | Induced CCR value ± Deviation | | | | |
| --- | --- | --- | --- | --- | --- |
|  | Glutamate | Malate | Proline+Pyruvate | Succinate | G3P |
| **LARVA** |  |  |  |  |  |
| **1 larva** | $1.5 \pm 0.4^a$ | $1.3 \pm 0.2^a$ | $1.4 \pm 0.1^a$ | $1.5 \pm 0.3^a$ | $1.1 \pm 0.1^b$ |
| **5 larvae** | $1.3 \pm 0.1^c$ | $1.2 \pm 0.1^c$ | $1.1 \pm 0.0^{cd}$ | $1.1 \pm 0.0^d$ | $1.2 \pm 0.1^c$ |
| **PUPA** |  |  |  |  |  |
| **1 pupa** | $1.4 \pm 0.2^e$ | $1.3 \pm 0.1^{ef}$ | $1.4 \pm 0.1^{ef}$ | $1.2 \pm 0.1^f$ | $1.6 \pm 0.2^g$ |
| **5 pupae** | $1.3 \pm 0.1^h$ | $1.3 \pm 0.1^h$ | $1.4 \pm 0.1^h$ | $1.1 \pm 0.1^i$ | $1.2 \pm 0.1^{hi}$ |
| **ADULT** |  |  |  |  |  |
| **1 thorax** | $1.1 \pm 0.3$ | $1.3 \pm 0.3$ | $1.3 \pm 0.3$ | $1.2 \pm 0.1$ | $1.1 \pm 0.1$ |
| **5 thoraces** | $1.8 \pm 0.6^j$ | $1.2 \pm 0.2^{jk}$ | $1.6 \pm 0.4^{jk}$ | $1.0 \pm 0.1^k$ | $1.1 \pm 0.1^k$ |

For permeabilized thoraces, the highest induced CCR values were observed with five permeabilized thoraces. In the case of a single permeabilized thorax, the highest RCC value was obtained for the substrates malate (1.3 ± 0.3) and proline+pyruvate (1.3 ± 0.3), with no significant differences found between substrates (ANOVA: F= 1.9131, df=4, p≥0.005). For five permeabilized thoraces, the highest calculated induced CCR value was found with the glutamate substrate (1.8 ± 0.6), and significant statistical differences were found between glutamate-succinate and glutamate-G3P substrates (KW test H: (4, N=38) = 17.59287 p≤0.05) (Table 2).

### Comparison of respiratory parameters of isolated mitochondria vs. permeabilized tissue

Respiratory parameters were compared between isolated mitochondria and permeabilized individuals at stage 3 and for each substrate across the developmental stage of *Ae. aegypti*. In the larval stage, higher oxygen consumption was obtained with the substrate glutamate and malate by L3-L4 larvae, which decreased slightly in pupae and rapidly in adults. With proline + pyruvate, the values recorded in L3-L4 larvae increased slightly in pupae and decreased in adults. For succinate, isolated mitochondria showed a progressive increase in oxygen consumption larvae L3-L4 to adults. However, this behavior was not observed with five permeabilized individuals, where the values decrease to nearly half of those recorded for larvae and pupa. For G3P, high oxygen consumption was obtained in L3-L4 larvae, which decreased for pupae. In adults, two different trends were noted: oxygen consumption increased in isolated mitochondria, while for permeabilized thoraces were similar to those recorded for pupae (Figure 5).

**Figure 5.**
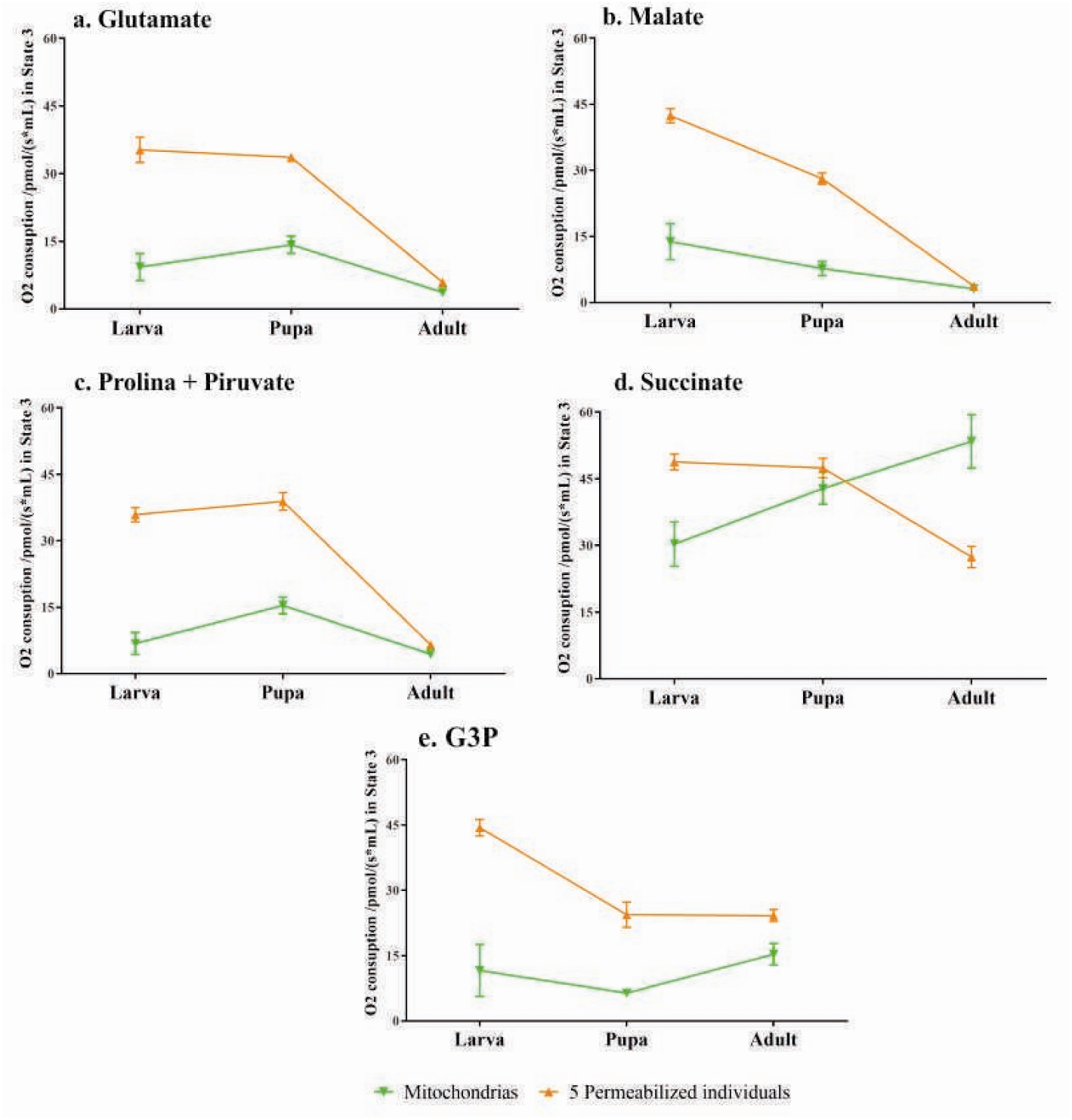
Average oxygen consumption (pmol/(sec*mL)) in state 3 was measured for isolated mitochondria and groups of five and four permeabilized individuals from, pupae, and adult females (thorax) of *Ae. aegypti* with each of the oxidizable substrates evaluated: glutamate, malate, proline+pyruvate, succinate, and G3P.

State 2 measurement was performed to quantify and compare the values of oxygen consumption across all respiratory states for isolated mitochondria and permeabilized individuals using each oxidizable substrate throughout the developmental stages. Oxygen consumption in state 2 (basal respiration) provides insight into the presence of potential endogenous substrates that contribute electrons to the mitochondrial respiratory chain. It is noteworthy that the values obtained in stage 2 are independent of the substrate used. A higher initial respiratory rate was observed in L3-L4 larvae, which progressively decreased to values below 8 pmol/(sec*mL) in the adult stage both in isolated mitochondria and in permeabilized tissue (Figure 6).

**Figure 6.**
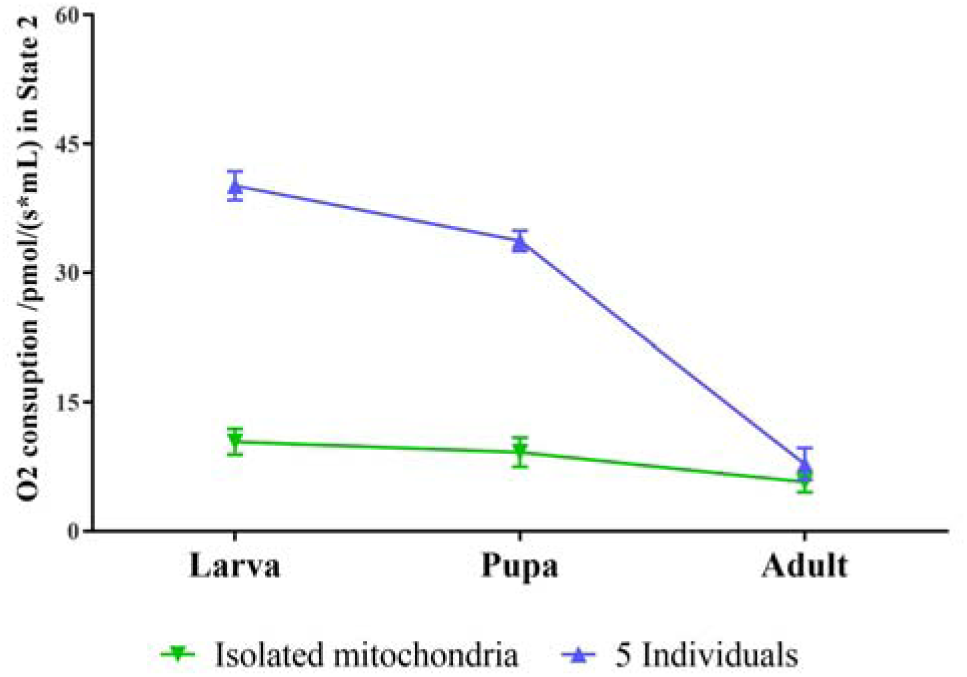
Average oxygen consumption (pmol/(sec*mL)) at stage 2 for isolated mitochondria and permeabilized individuals (5) of larvae (L4), pupae, and adult females (thorax) of *Ae. aegypti*.

### Enzyme activity

To understand the functioning of the mitochondrial respiratory chain in each of the developmental stages of *Ae. aegypti*, the activity of eight enzymes associated with the four respiratory complexes and the ATPase was evaluated.

#### Complex I

The activity of Complex I was evaluated by quantifying the entry of electrons from NADH (NADH oxidase enzyme) and their subsequent passage through the complex (NADH dehydrogenase enzyme). A higher electron input (NADH oxidase activity) was found in mitochondria isolated from thorax of adult females, with measurement of 40.3 ± 5.8 pmol O_2_/ sec*mL protein, followed by pupae at 15.3 ± 1.7 pmol O2/ sec*mL protein and larvae 8.6 ± 0.6 pmol O2/ sec*mL protein, with significant differences between the stages evaluated (ANOVA: F= 433.089, df=2, p=0.000124) (Figure 7a). The high activity of NADH oxidase in adults is associated with the oxidation of substrates via complex I, while this enzymés activity progressively decreases in the larval stage, with lower activity values.

**Figure 7.**
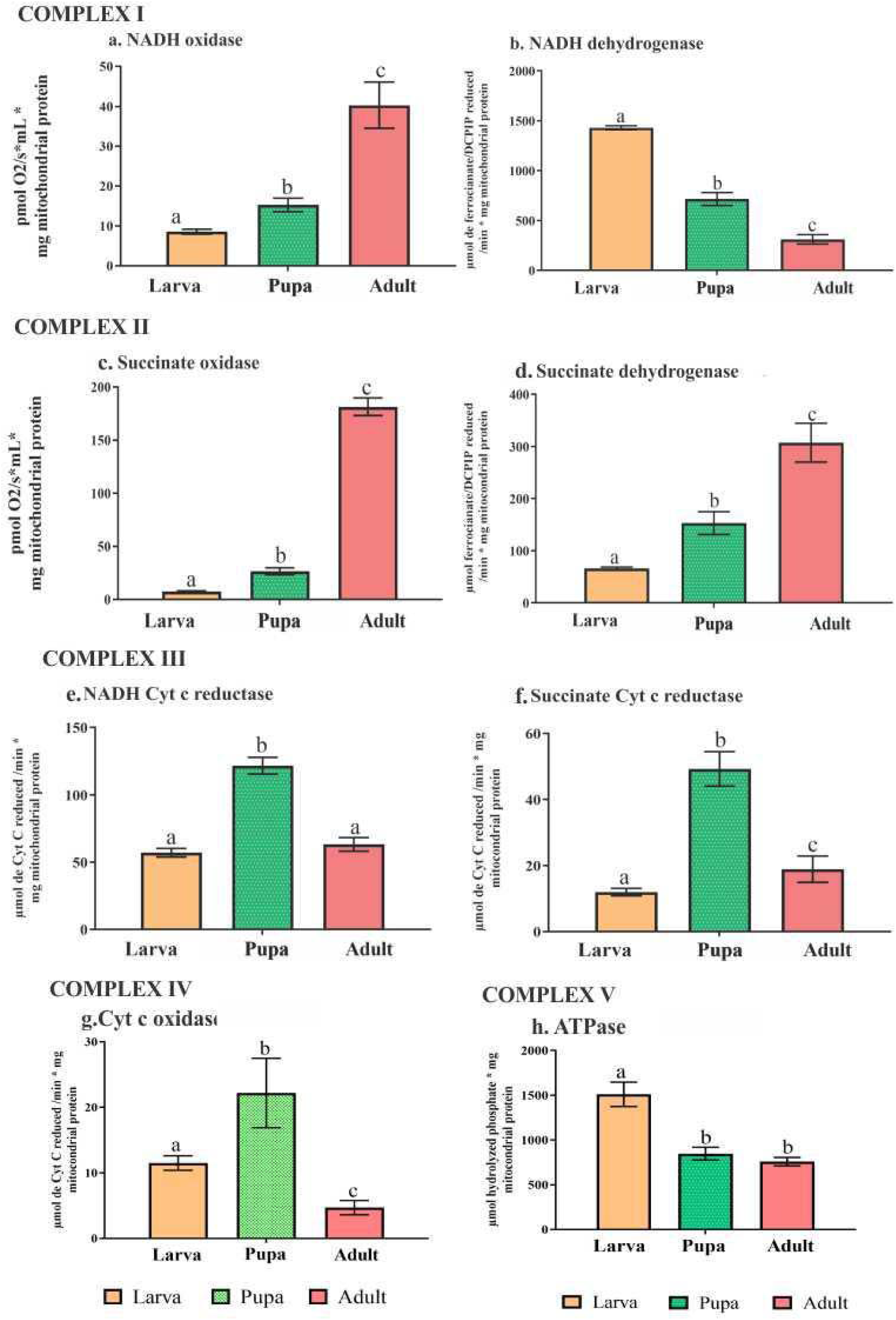
Measurement of CTE specific enzyme activity for mitochondria isolated from L4 larvae, pupae, and thorax of *Ae. aegypti* females. Different letters indicate significant statistical differences between enzyme activity for each developmental stage. a. NADH oxidase a,b,c (ANOVA test p=0.00124); b. NADH dehydrogenase a,b,c (ANOVA test p=0.000125); c. Succinate oxidase a,b,c (ANOVA test p=0.00120); d. Succinate dehydrogenase a,b,c (ANOVA test p=0.000127); e. NADH Cyt c reductase a,b (ANOVA test p=0.00168); f. Succinate Cyt c reductase a,b,c (ANOVA test p=0.000244); g. Cytochrome oxidase a,b,c (ANOVA test p=0.000212) and h. ATPase a,b (ANOVA test p=0.000144).

For NADH dehydrogenase, the larval stage exhibited the highest activity, with 1430.5 ± 21 μmol of reduced ferrocyanate/min*mg of mitochondrial protein, indicating an increased electron transfer through this complex. The pupal stage followed, with 715 ± 64.3 μmol of reduced ferrocyanate/min* mg of mitochondrial protein, while the lowest activity was observed in adults, with 310.2 ± 47.2 μmol of reduced ferrocyanate/min*mg of mitochondrial protein. Statistical analysis revealed significant differences between developmental stages (ANOVA test F= 1954.67, df=2, p=0.000125) (Figure 7b). The higher activity recorded in larvae suggests a higher electron influx when using glutamate as a substrate, as well as with the proline+pyruvate. Although pupae do not show marked activity, their observed values denote efficient electron entry and passage from NADH.

#### Complex II

Complex II was assessed by quantification of the enzymes succinate oxidase and succinate dehydrogenase, which quantify the entry of electrons from succinate and their passage through complex II, respectively.

For succinate oxidase, a higher electron input was observed for mitochondria isolated from the thorax of adult females with 181.5 ± 8.3 pmol of O2/ sec*mL of protein, followed by the activity recorded for pupae 26.7 ± 3.2 pmol of O2/ sec*mL of protein and, finally, the activity of L4 larvae 7.5 ± 0.6 pmol of O2/ sec*mL of protein, with significant differences between these groups (ANOVA: F= 8233.60, df=2, p=0.00120) (Figure 7c).

Consistent with higher electron influx via Succinate, the adult stage presented the highest succinate dehydrogenase enzyme activity (307.4 ± 37.2 μmol of reduced DCPIP/min*mg of mitochondrial protein), followed by the pupal stage (152, 9 ± 22 μmol of reduced DCPIP/min*mg of mitochondrial protein) and lower activity in the larval stage (66 ± 2.7 μmol of reduced DCPIP/min*mg of mitochondrial protein), with significant differences between them (ANOVA: F= 245.140, df=2, p=0.000127) (Figure 7d).

For both enzymes evaluated, the adult stage presented the highest activity values, demonstrating an effective electron transfer from complex II to complexes III and IV. This indicates a greater electron flow in adults compared to the juvenile stages (larva and pupa). Notably, succinate oxidase activity was very low in larvae and pupae, suggesting that electron entry into respiratory chain does not primarily via complex II in these stages.

#### Complex III

Complex III was evaluated by quantifying electrons transfer from NADH through Complex I to Cytochrome c (Cyt c) (NADH Cyt c reductase enzyme) and by quantifying the passage of electrons from succinate via complex II through FAD, passing through the Fe-S centers and subsequently to the electron acceptor DCPIP (succinate Cyt c reductase enzyme).

The pupal stage exhibited the highest activity for the NADH Cyt c reductase (121.6 ± 6.2 μmol of Cyt c reduced/min*mg of mitochondrial protein), followed by adult stage (63.2 ± 5, 1 μmol of Cyt c reduced/min*mg mitochondrial protein) and larva (57 ± 3.2 μmol of Cyt c reduced/min*mg mitochondrial protein), with significant differences between pupa and larva-adult stage values (ANOVA: F= 329.244, df=2, p=0.00168) (Figure 7e).

When quantifying Succinate Cyt c reductase activity, the pupal stage exhibited the highest activity (49.3 ± 5.2 μmol of Cyt c reduced/min*mg of mitochondrial protein), followed by adult stage (18.9 ± 4 μmol of Cyt c reduced/min*mg of mitochondrial protein) and larva stage (12 ± 1.1 μmol of Cyt c reduced/min*mg of mitochondrial protein), with significant differences between the stages evaluated (ANOVA: F= 59.8782, df=2, p=0.000244) (Figure 7f).

The marked activity of NADH and Succinate Cyt c reductase in the pupal stage represents a novel finding, likely related to the fact that the pupa does not feed. This, suggests that pupa must rely on products of glycolysis and the Krebs cycle intermediates, such as succinate, to supply electrons to the mitochondrial respiratory chain.

#### Complex IV

Complex IV was evaluated by monitoring the oxidation of Cyt c, which transfers one electron at a time to complex IV of the respiratory chain.

The pupal stage presented the highest activity with 22.2 ± 5.3 μmol of Cyt c reduced/min*mg of mitochondrial protein, followed by the larval stage 11.5 ± 1.1 μmol of Cyt c reduced/min*mg of mitochondrial protein, and finally the adult stage 4.7 ± 1.1 μmol of Cyt c reduced/min*mg of mitochondrial protein with significant differences between the stages evaluated (ANOVA: F= 52.1293, df=2, p=0.000212) (Figure 7g).

The higher of Cyt c oxidation in the pupae stage is directly related to the activity recorded for cytochrome c reductases (Complex III), facilitating the passage and efficient utilization of electrons derived from the oxidation of substrates via complexes I and II. In contrast, activity values of adult individuals were lower, suggesting a possible regulatory mechanism for membrane potential generation and increased oxidative stress. Considering the enzyme complexes results described above, it is evident that oxidase activity in adults is directly linked to the activity of complexes II and III of the respiratory chain.

#### Complex V

ATPase activity was evaluated as an enzyme with reversible functionality, meaning its activity is directly associated with ATP synthase funtion. The higher activity of this enzyme was observed in mitochondria isolated from L4 larvae (1510.9 ± 135.8 μmol of hydrolyzed phosphate*mg of mitochondrial protein), followed by pupal stage (846, 8 ± 71 μmol of hydrolyzed phosphate*mg of mitochondrial protein), and finally, the adult stage (757.5 ± 47.1 μmol of hydrolyzed phosphate*mg of mitochondrial protein) finding significant differences between larval and pupal-adult values (ANOVA: F= 148.872, df=2, p=0.000144). The decrease in enzyme activity throughout mosquito development may be directly related to stage-specific metabolic and physiological processes in turn influence the mitochondrial energy demand (Figure 7h).

## Discussion

### Respiratory parameters in isolated mitochondria

The mosquito *Ae. aegypti* is a holometabolous insect, a characteristic that results in drastic metamorphic changes throughout its life cycle, modifying various metabolic functions. These changes are represented in fluctuations in cellular energy and oxygen demands (Merkey et al. 2011). One of the most widely used methods to assess physiological alterations in *Ae. aegypti* and other insects involves measuring mitochondrial bioenergetic parameters, such as oxygen consumption and enzymatic activity of the electron transport chain under different conditions (Correa et al., 2015).

In this study, we describe for the first time the mitochondrial respiratory behavior of *Ae. aegypti* throughout its life cycle o in response to different oxidizable substrates. The highest oxygen consumption values were observed when using succinate as a substrate, starting at the larval stage (L4) and progressively increasing until the adulthood (Supplementary material - Figure 1). This result would indicate an increased Complex II activity, mediated by the oxidation of succinate to fumarate via Succinate dehydrogenase (Mailloux, 2015; Moosavi et al., 2019).

In larval stages, high oxygen consumption values were observed when using succinate as a substrate, compared to the values obtained with other evaluated substrates (Figure 1). These values may be associated with the need for biomass acquisition and energy storage.

Although no studies have directly examined mitochondrial activity in *Aedes aegypti* larvae, Jacobs et al. (2020) provide insights into the role of mitochondrial metabolism in regulating larval growth in insects, specifically in the model organism *Drosophila melanogaster*. The authors state that respiratory chain activity and oxidative phosphorylation can be physiologically regulated by nutrient availability in the medium, the potential presence of xenobiotics, and environmental disturbances.

Considering that the larval stage is primarily dedicated to biomass accumulation for later ATP production and biomolecule synthesis, dietary differences and/or deficiencies could significantly alter the individual’s normal physiological function.

Considering the biomass acquisition requirements during the larval stages, succinate oxidation via complex II is particularly relevant In the absence of proton entry through the mitochondrial membrane, the potential gradient is lower, leading to reduce reactive oxygen species (ROS) generation (Cecchini, 2003). This aspect is important because L4 larvae do not feed during the middle of their instar in preparation for the physiological changes characteristic of the pupal stage, so there is less energy expenditure in detoxification mechanisms (Bounias et al., 1989; Merkey et al., 2011; Jacobs et al., 2020).

It is worth noting that, although oxygen consumption with the other substrates was significantly lower, NADH molecules via Complex I (derived from the metabolism of glutamate, malate, and proline + pyruvate) make an important contribution to the proton gradient. Their role in the electron transport chain is essential for maintaining proper mitochondrial function (Figure 7). Additionally, the electron supply from these three substrates may be directly linked to energy reserves in insects. Unlike other animals, insects do not have large glycogen reserves; instead, trehalose serves as the primary sugar source in the hemolymph. The limited glycogen stores found in insects are primarily stored in fat body tissue and the intestine, fluctuating in response to motor activity, environmental conditions, and becoming nearly depleted after exoskeleton molting periods. (Thompson, 2003).

The pupal stage is a non-feeding phase characterized by extensive tissue reorganization for the development of adult organs. This metabolic shift leads to a decline in biomass production, as stored lipids are metabolized to sustain cell division, tissue differentiation, and hatching (Jacobs et al., 2020). According to Jacobs et al. (2020), mitochondrial activity during this stage in *Drosophila melanogaster* is lower than in larval and adult stages, following a U-shaped metabolic curve (Merkey et al., 2011). However, a key distinction is that *Drosophila* undergoes a completely terrestrial life cycle, with pupae remaining immobile until emergence. In contrast, *Aedes aegypti* juveniles are aquatic, and while pupae are less mobile than larvae, they can still respond to external stimuli.

Despite their limited mobility, pupae exhibit intense metabolic activity, which is essential for adult tissue formation. This increased energy demand is met through glycolysis and the Krebs cycle, utilizing succinate and coenzyme Q, which transfers electrons from NADH and FADHLJ oxidation to Complex III. Additionally, coenzyme Q plays a crucial role in preventing lipid peroxidation, acting as an antioxidant system (Deshwal et al., 2023). This metabolic adaptation may explain the higher oxygen consumption values observed when using succinate (via Complex II), as well as glutamate and proline + pyruvate (via Complex I). Furthermore, the marked activity of NADH– cytochrome c reductase and succinate–cytochrome c reductase supports this enhanced mitochondrial function.

The adult phase is the only stage in the *Aedes aegypti* life cycle for which mitochondrial physiology has been more extensively characterized. Most published studies on this topic focus on the mitochondrial bioenergetics of adult individuals, typically isolating mitochondria from flight muscles due to their larger mitochondrial size and high respiratory activity rates (Correa et al., 2015; Gonçalves et al., 2009). Several studies highlight the key role of proline as an energy source in the flight muscles of *Ae. aegypti* (Goldstrohm et al., 2003; Scaraffia & Wells, 2003). The role of this amino acid as an oxidizable substrate in mosquito mitochondria has been further detailed in the work of Correa et al. (2015) and Gonçalves et al. (2009), who describe proline + pyruvateand glycerol 3-phosphate (G3P) as the primary substrates contributing to ATP formation in adult males and females. It is important to note that these authors identified small metabolic differences between adults based on dietary sources. In females, blood feeding provides proline, whereas in males, sugar feeding supplies glucose. Electron input occurs primarily via Complex I or through the direct electron transfer to ubiquinone, facilitated by specific dehydrogenases, including proline dehydrogenase (ProDH) and G3P dehydrogenase (G3PDH) (Correa et al., 2015; Gonçalves et al., 2009).

When comparing the oxygen consumption values in stage 3 with the proline + pyruvate substrate reported by Goncalves et al., (2009) (43.47±5.09 nmol/(min*mg)) and Correa et al., (2015) (132 ± 58 nmol/(min*mg)) in sugar-fed females, the values from these previous studies are more than twice those values reported in the present study (4.4 ± 0.9 pmol/(s*mL)). Although these authors noted that respiration rates with this substrate are low compared to values obtained to other insects, as their primary energy source is fatty acid oxidation (Correa et al., 2015); the results obtained in this study are paradoxically even lower than those previously reported.

A similar trend was observed for G3P oxidation, where respiratory values reported by previous studies (Gonçalves et al., 2009: 179.9 ± 35.78 nmol/(min·mg); Correa et al., 2015: 68 ± 28 nmol/(min·mg)) were notably higher than those obtained in this study (15.3 ± 2.5 pmol/(s·mL)). Correa et al. (2015) highlighted that, given the high metabolic activity of flight muscles, increased respiratory rates with proline + pyruvate suggest complete depletion of glycogen stored in fat bodies and muscle tissue. This oxidation pathway is particularly active in females, as they acquire proline directly from the blood diet. However, newly emerged females do not seek blood meals immediately after eclosion; instead, they first focus on carbohydrate ingestion to rapidly replenish glycogen reserves. This metabolic shift relies on larval biosynthetic reserves (stored in fat tissue and the intestine), which limits flight potential and survival time until high-sugar food sources are accessed (Briegel et al., 2001; Dou et al., 2023).

Since the oxygen consumption values in this study differed significantly from those reported by Gonçalves et al. (2009), we sought to confirm the mitochondrial integrity of L4 larvae using the Borrero et al. (2018) protocol, by calculating the Respiratory Control Coefficient (RCC). Although it cannot be definitively concluded that the mitochondria isolated in this study were uncoupled, the lower-than-expected values may be associated with mechanical or chemical damage to the outer membrane during the isolation process (Hartwig et al., 2015). This is consistent with previous findings that mitochondrial isolation alters functional interactions between mitochondria and the extramitochondrial environment (Saks et al., 1998). Despite our effort, we were unable to replicate the RCC values reported in the literature, even when using permeabilized tissue at all developmental stages,where mechanical damage from isolation would not be expected This suggest that t baseline RCC values may vary between the laboratory colonies studied.

### Measurement of respiration in permeabilized larvae, pupae, and female thoraxes

An essential aspect of mitochondrial physiology studies is the preservation of cellular architecture and interactions between organelles in situ (Saks et al., 1998). To address this, various authors have proposed methods to study mitochondria in vivo without the need for tissue isolation (Beenakkers et al., 1984; Candy, 1970; Saks et al., 1991). The tissue permeabilization technique allows the evaluation of mitochondrial function in vivo while avoiding mechanical or chemical procedures that could induce structural and functional alterations (Beenakkers et al., 1984; Saks et al., 1991).

Considering the advantages of using permeabilized tissue and to minimize potential mitochondrial damage associated with isolation procedures, respiratory parameters were evaluated in permeabilized tissues of larvae, pupae, and thorax of *Ae. aegypti* adults. Considering the absence of studies on permeabilized tissue in juvenile stages, a comparison was performed using 1 and 5 individuals to determine the minimum number of specimens required for this type of experiment. Overall, the respiratory values obtained with permeabilized tissue were comparable to those observed in isolated mitochondria. When using succinate as a substrate, the highest oxygen consumption values were recorded across all developmental stages.However, this result does not imply that Complex II is the sole pathway for electron entry into the mitochondrial respiratory chain. Instead, the findings suggest a substrate preference that varies according to developmental stage.

Oxygen consumption in stage 2 enables the quantification of potential endogenous substrates that supply electrons to the mitochondrial respiratory chain (Nicholls et al., 2012). The presence of these substrates was higher in permeabilized individuals, particularly in juvenile stages (larva and pupa). In larva L4, high respiratory values were observed at stage 2, especially in permeabilized larvae (40.1 ± 1.7 nmol/(min*mg)), indicating endogenous substrates consumption prior to the addition of exogenous substrates. The highest oxygen consumption was recorded with succinate as the substrate. Although statistically significant differences were found among evaluated substrates, the recorded oxygen consumption and calculated bioenergetic capacity values suggest additional electron input via Complex I (glutamate) and via glycerol 3-phosphate dehydrogenase (G3PDH).

Juvenile individuals of the family Culicidae are characterized by aquatic habits and *Ae. aegypti* larvaepossess specialized structures, such as the respiratory siphon, which allow them to obtain oxygen directly from the air while remaining near the water surface. During their aquatic life, the juvenile stages (larvae and pupae) frequently experience periods of hypoxia, during which they descend to the bottom to feed and evade potential threats. Before reaching a potentially anoxic state, these individuals must restore their metabolic status and replenish the necessary energy in their tissues (Redecker & Zebe, 1988).

One adaptive strategy that provides flexibility in responding to rapid environmental changes involves activating the fatty acid oxidation pathway in the Krebs cycle to generate succinate. It has been reported that when an active tissue does not receive sufficient oxygen to meet its metabolic demands, succinate accumulates, playing a crucial role in the tissue re-oxidation process (Pekny et al., 2018). This mechanism presents a significant advantage: since succinate oxidation via Complex II does not involve proton pumping across the mitochondrial membrane, it helps maintain mitochondrial redox balance, reduces electron availability for reactive oxygen species (ROS) formation, and prevents a decline in ATP production (Bringaud et al., 2006).

Although endogenous substrate consumption was also recorded in pupae, the values obtained were lower than those recorded for L4 larvae (isolated mitochondria: 9.2 ± 1.7 nmol/(min*mg); permeabilized pupae 33.7 ± 1.2 nmol/(min*mg)). For most of the evaluated substrates, oxygen consumption in pupae was slightly lower than in larvae, with a higher respiratory rate observed with succinate as the substrate. Unlike larvae, no significant statistical differences were found between succinate and proline+pyruvate combination, indicating that respiration in this stage occurs through the entry of electrons via Complex II and Ubiquinone (Proline dehydrogenase - PDH).

As in the larval stage, pupa also undergoes periods of hypoxia, activating the fatty acid oxidation pathway to generate succinate as an electron source and to restore oxidative capacity (Pekny et al. 2018). Although proline levels in *Ae. aegypti* larvae and pupae are low (Goldstrohm et al., 2003), this amino acid plays a crucial role in metabolism and cuticle synthesis during molting, serving as an energy supplement (Bounias et al., 1989; Chaput & Liles, 1969). Most holometabolous insects accumulate large lipids reserves until the middle of their last larval instar (at which point feeding ceases, and these reserves are utilized for energy during pupation e (Beenakkers et al., 1981).Therefore, the observed decrease in oxygen consumption values is likely associated with mitochondrial mediation of cell death processes required for tissue remodeling, leading to reduced energy demand and decreased active ion transport (VenkatRao et al., 2016).

In adults, a low state 2 mitochondrial respiration rate was observed, indicating minimal consumption of endogenous substrates. This result is consistent with the findings of Correa et al. (2015), who evaluated the mitochondrial physiology of *Aedes aegypti* in permeabilized male and female thoraxes. Regarding state 3 oxygen consumption, our study found the highest consumption rate with succinate as the substrate, followed by glycerol-3-phosphate (G3P), with no statistically significant differences between them. However, Correa et al. (2015) reported the highest oxygen consumption with proline+pyruvate and G3P.

These authors described that approximately 90% of the electrons used in oxidative phosphorylation in flight muscles originate from proline+pyruvate metabolism. Part of these electrons are derived from proline oxidation via four pathways: direct transfer to ubiquinone by proline dehydrogenase (PDH), the action of carboxylate dehydrogenase, glutamate dehydrogenase, and enzymes of the Krebs cycle. Additionally, the high respiratory rates observed with this substrate explain the total depletion of glycogen stored in fat bodies and thoracic muscles after flight periods (Correa et al., 2015).

Goldstrohm et al. (2003) also reported high proline levels in unfed females, which increased significantly after blood feeding. Moreover, they observed that in males, proline levels also rose following sugar feeding, suggesting that proline serves as a temporary storage form of toxic ammonium, released during the degradation of dietary amino acids. This ammonium is stored as proline and ultimately excreted as urea or uric acid.

In this study, the oxygen consumption values with the G3P substrate in adults were again lower than those reported by Correa et al. (2015). G3P production is associated with the activity of glycerol kinase during lipolysis in flight muscle. The enzyme allows the utilization of G3P to meet the high energy demand of muscle (Beenakkers et al. 1981). In permeabilized tissue, the cytosolic form of glycerol 3-phosphate dehydrogenase (G3PDH) may predominate, which is responsible for the reoxidation of extra-mitochondrial NADH derived from glycolysis and increasing electron input to oxidative phosphorylation (Correa et al. 2015). Although this increased electron input could be directly related to hydrogen peroxide (H2O2) production, mitochondria can regulate electron input by quantifying ADP and ATP present in the cell as mechanisms controlling G3PDH and glycerol phosphate shuttle activity (Gaviraghi & Oliveira, 2019). Additionally, it has been reported that part of G3P produced in flight muscle is released directly into the hemolymph for transport to the fat body (Beenakkers et al. 1981), so that part of this substrate would not enter oxidative phosphorylation.

Similar to the experiments performed with isolated mitochondria, the CCR was calculated in permeabilized tissue. When comparing the results obtained from isolated mitochondria and permeabilized tissue, The use of five permeabilized individuals was deemed optimal for measuring mitochondrial physiological parameters. Furthermore, since our respiration results in larvae, pupae and adults were consistent between extracted mitochondria and permeabilized tissue, we concur with Correa et al. (2015). These authors previously concluded that, given the similarity in oxygen consumption values between isolated mitochondria and permeabilized tissue, mitochondrial physiological parameters can be assessed in situ without the need to perform an isolation protocol.

### Enzyme activity

This is the first study to describe the activity of enzymes related to the mitochondrial respiratory chain at different stages of the *Ae. aegypti* life cycle, without focusing on enzyme activity in response to substances of natural or synthetic origin. In the larval stage, intrinsic activity of complexes I and II was observed, likely associated with the consumption of endogenous substrates, as described above. The activity of oxidases suggests that electron entry occurs similarly with glutamate and succinate as substrates, although with greater efficiency via Complex I (Glutamate). These findings are consistent with the oxygen consumption values obtained with permeabilized larval tissue, where electron entry via Complex I and II was recorded.

Furthermore, a higher electron flux through Complex was observed based on the activity of NADH Cyt c reductase and Cytochrome c oxidase. Consistent with the effective passage of electrons via Complex I to Complex IV, high ATPase enzyme activity was recorded, possibly related to a higher energy requirement (ATP production) due to the synthesis of energy storage tissues (lipids) and the high metabolic activity of the larva (Jacobs et al., 2020; Merkey et al., 2011).

For pupae, higher activity of complex II-related enzymes was recorded, both in the oxidative activity of the enzyme succinate oxidase and in the intrinsic activity of the complex (succinate dehydrogenase). This increased activity supports the oxygen consumption values recorded with the succinate substrate, which, as mentioned above, serves as an electron source after periods of hypoxia. Additionally, for the pupal stage, a high activity of reductase enzymes was observed, particularly with NADH Cyt c, denoting an effective passage of electrons to Complex III, primarily from Ubiquinone. Although this result may initially appear contradictory to the oxidases activity present in this study, it should be considered that the use of the substrate succinate possibly occurs only when the tissues need to reactivate oxidative phosphorylation after a period with low oxygen, therefore, the entry of electrons with the substrate proline+pyruvate contributes significantly to the activity recorded. The high activity of Cyt c oxidase may be related to the underlying cell death processes (VenkatRao et al., 2016), being directly related to the low ATPase activity and low substrate concentration (NADH and succinate), since pupae must maintain a slow basal metabolism to use as many energy sources (fatty acids) as possible for the processes of histolysis and histogenesis in the development of the adult individual (Beenakkers et al. 1981).

For adults, higher succinate oxidase enzyme activity was observed, consistent with the results obtained for oxygen consumption in the present study. NADH oxidase activity was also detected, indicating substrate oxidation via Complex I. Upon evaluating the intrinsic activity of dehydrogenases in complexes I and II, a higher succinate dehydrogenase activity was recorded. When analyzing electrons transfer to complex III, this process was found to be primarily mediated by the enzymes of complexes I and II, with a slight increase in the activity of the enzyme NADH Cyt c reductase (Complex I - glutamate and proline+pyruvate). Consistent with the reported by Correa et al. (2015), this study revealed low cyt c oxidase activity, suggesting a potential regulation of membrane potential generation, possibly due to a possible oxidative stress of these mitochondria.

In general terms, significant changes in the enzymatic function of the electron transport chain occur throughout the life cycle of the *Ae. aegypti*. During the larval phase, there is a substantial influx of electrons via complexes I and II, leading to increased ATP production. This energy is essential for tissue formation, molecule synthesis, energy expenditure during the four larval stages, and glycogen storage for subsequent developmental stages. This activity declines during the pupal stage, which relies mainly on lipid reserves and glycogen stored in the fat body as the main energy source.

Although low ATPase activity was found in adults, energy requirements are modulated by various factors such as the type of feeding, availability of food rich in carbohydrates and sugars before the first blood feeding in females, availability of sugars (glucose) in males, availability and use of amino acids (proline) in females during gonadotropic cycles and energy expenditure during flight.

### Conclusions

Although mitochondrial isolates remain the most widely used method to assess mitochondrial function in insects, permeabilized tissue allows in situ measurement of these parameters without altering mitochondrial biochemistry. This approach reduces the number of individuals required per developmental stage while preserving cellular interactions and physiological relevance.

This study provides the first description of mitochondrial respiratory behavior throughout the life cycle of *Aedes aegypti*using both isolated mitochondria and permeabilized tissue. The highest oxygen consumption (state 3) was observed with succinate as the primary oxidizable substrate across all developmental stages, with a progressive increase from larvae to adults. These results align with enzymatic activity data, showing elevated succinate oxidase and succinate dehydrogenase (Complex II) activity.

Mitochondrial respiration in larvae is driven by electron input via succinate (Complex II) and substrates from Complex I (glutamate, malate, and proline+pyruvate), supporting glycogen storage. In pupae, respiration primarily relies on Complex II and proline dehydrogenase, with a shift towards fatty acid oxidation during hypoxic periods to generate succinate.

In sugar-fed females, respiration is dominated by electron flux through Complex II, with high succinate oxidase activity suggesting a role in ROS regulation, particularly in flight muscle. G3P utilization, linked to glycerol kinase activity during lipolysis, helps sustain energy demands by depleting glycogen reserves in fat bodies and thoracic muscles.

## Acknowledgments

To the Centro de Investigaciones en Enfermedades Tropicales CINTROP, through the Grupo de Entomología Medica; the Grupo de Investigación en Bioquímica y Microbiología-GIBIM; Proyecto Biorreto “Desarrollo de nuevos productos con actividades antimicrobiana, insecticida y repelente a partir de moléculas aisladas de plantas aromáticas colombianas” (Development of new products with antimicrobial, insecticidal and repellent activities from molecules isolated from Colombian aromatic plants). To Luis Carlos Vesga for the review and comments to the final document.

## Author contributions

RMCM and DLUD contributed substantially to the design of the work, data acquisition, data analysis and writing of the article; SCMS and JED contributed to the study design, analysis, interpretation of data, and writing of the article.

## Notes

### Competing Interest Statement

The authors have declared no competing interest.

## References

Bajda, S., Dermauw, W., Panteleri, R., Sugimoto, N., Douris, V., Tirry, L., Osakabe, M., Vontas, J., Van Leeuwen, T., 2017. A mutation in the PSST homologue of complex I (NADH: ubiquinone oxidoreductase) from Tetranychus urticae is associated with resistance to METI acaricides. Insect Biochem. Mol. Biol. 80, 79–90. 10.1016/j.ibmb.2016.11.010

Beenakkers, A.M.T., Van der Horst, D.J., Van Marrewijk, W.J.A., 1984. Insect flight muscle metabolism. Insect Biochem. 14, 243–260. 10.1016/0020-1790(84)90057-X

Beenakkers, A.M.T., Van der Horst, D.J., Van Marrewijk, W.J.A., 1981. Role of Lipids in Energy Metabolism, in: Downer, R.G.H. (Ed.), Energy Metabolism in Insects. Springer US, Boston, MA, pp. 53–100. 10.1007/978-1-4615-9221-1_3

Bhatt, S., Gething, P., Brady, O., Messina, J., Farlow, A., Moyes, C., Drake, J., Brownstein, J., Hoen, A., Sankoh, O., Myers, M., George, D., Jaenisch, T., Wint, W., Simmons, C., Scott, T., Farrar, J., Hay, S., 2013. The global distribution and burden of dengue. Nature 496, 504–507. 10.1038/nature12060

Bolter, C.J., Chefurka, W., 1990. Extramitochondrial release of hydrogen peroxide from insect and mouse liver mitochondria using the respiratory inhibitors phosphine, myxothiazol, and antimycin and spectral analysis of inhibited cytochromes. Arch. Biochem. Biophys. 278, 65–72. 10.1016/0003-9861(90)90232-N

Borrero-Landazabal, M.A., Duque, J.E., Mendez-Sanchez, S.C., 2020. Model to design insecticides against *Aedes aegypti* using in silico and in vivo analysis of different pharmacological targets. Comp. Biochem. Physiol. C. Toxicol. Pharmacol. 229, 108664. 10.1016/j.cbpc.2019.108664

Borrero, M.A., Carreño, A.L., Kouznetsov, V.V., Duque Luna, J.E., Mendez-Sanchez, S.C., 2018. Alterations of mitochondrial electron transport chain and oxidative stress induced by alkaloid-like α-aminonitriles on *Aedes aegypti* larvae. Pestic. Biochem. Physiol. 144, 19–26. 10.1016/j.pestbp.2017.11.006

Bounias, M., Vivarès, C.P., Nizeyimana, B., 1989. Functional relationships between free amino acids in the hemolymph of fourth instar larvae of the mosquito *Aedes aegypti* (Diptera, Culicidae) as a basis for toxicological studies. J. Invertebr. Pathol. 54, 16–22. 10.1016/0022-2011(89)90133-x

Bradford, M.M., 1976. A rapid and sensitive method for the quantitation of microgram quantities of protein utilizing the principle of protein-dye binding. Anal. Biochem. 72, 248–254. 10.1016/0003-2697(76)90527-3

Briegel, H., Knüsel, I., & Timmermann, S. E. 2001. *Aedes aegypti:* size, reserves, survival, and flight potential. Journal of Vector Ecology: Journal of the Society for Vector Ecology, 26(1), 21–31. https://www.ncbi.nlm.nih.gov/pubmed/11469181

Bringaud, F., Rivière, L., Coustou, V., 2006. Energy metabolism of trypanosomatids: adaptation to available carbon sources. Mol. Biochem. Parasitol. 149, 1–9. 10.1016/j.molbiopara.2006.03.017

Candy, D.J., 1970. Metabolic studies on locust flight muscle using a new perfusion technique. J. Insect Physiol. 16, 531–543. 10.1016/0022-1910(70)90192-7

Castillo-Morales, R.M., Carreño Otero, A.L., Mendez-Sanchez, S.C., Da Silva, M.A.N., Stashenko, E.E., Duque, J.E., 2019. Mitochondrial affectation, DNA damage and AChE inhibition induced by Salvia officinalis essential oil on Aedes aegypti larvae. Comparative Biochemistry and Physiology Part - C: Toxicology and Pharmacology 221. 10.1016/j.cbpc.2019.03.006

Castrillón, J., Carlos, J., Urcuqui, S., 2014. Dengue en Colombia, diez años de evolución. Revista Chilena de Infectología 32, 22–29. 10.4067/S0716-10182015000300002

Cecchini, G., 2003. Function and structure of complex II of the respiratory chain. Annu. Rev. Biochem. 72, 77–109. 10.1146/annurev.biochem.72.121801.161700

Chaput, R.L., Liles, J.N., 1969. Free and peptide-bound amino acids during the larval and pupal stages of the yellow-fever mosquito, *Aedes aegypti*. Ann. Entomol. Soc. Am. 62, 742–747. 10.1093/aesa/62.4.742

Colinet, H., Renault, D., Roussel, D., 2017. Cold acclimation allows Drosophila flies to maintain mitochondrial functioning under cold stress. Insect Biochem. Mol. Biol. 80, 52–60. 10.1016/j.ibmb.2016.11.007

Conde, A., 2003. Estudio de la longevidad y el ciclo gonotrofico del *Aedes aegypti*, Cepa Girardot en condiciones de laboratorio.

Correa, C.C., Aw, W.C., Melvin, R.G., Pichaud, N., Ballard, J.W.O., 2012. Mitochondrial DNA variants influence mitochondrial bioenergetics in Drosophila melanogaster. Mitochondrion 12, 459–464. 10.1016/j.mito.2012.06.005

Correa, J., Gaviraghi, A., Oliveira, M.F., 2015. Mitochondrial physiology in the major arbovirus vector *Aedes aegypti*: Substrate preferences and sexual differences define respiratory capacity and superoxide production. PLoS One 10, 1–35. 10.1371/journal.pone.0120600

Deshwal, S., Onishi, M., Tatsuta, T., Bartsch, T., Cors, E., Ried, K., Lemke, K., Nolte, H., Giavalisco, P., & Langer, T. 2023. Mitochondria regulate intracellular coenzyme Q transport and ferroptotic resistance via STARD7. Nature Cell Biology, 25(2), 246–257. 10.1038/s41556-022-01071-y

Dou, X., Chen, K., Brown, M. R., & Strand, M. R. 2023. Multiple endocrine factors regulate nutrient mobilization and storage in *Aedes aegypti* during a gonadotrophic cycle. Insect Science, 30(2), 425–442. 10.1111/1744-7917.13110

Gaviraghi, A., Oliveira, M.F., 2020. A simple and reliable method for longitudinal assessment of untethered mosquito induced flight activity. J. Insect Physiol. 126, 104098. 10.1016/j.jinsphys.2020.104098

Gaviraghi, A., Oliveira, M.F., 2019. A method for assessing mitochondrial physiology using mechanically permeabilized flight muscle of *Aedes aegypti* mosquitoes. Anal. Biochem. 576, 33–41. 10.1016/j.ab.2019.04.005

Giacomello, M., Pyakurel, A., Glytsou, C., Scorrano, L., 2020. The cell biology of mitochondrial membrane dynamics. Nat. Rev. Mol. Cell Biol. 21, 204–224. 10.1038/s41580-020-0210-7

Giulivi, C., Ross-Inta, C., Horton, A.A., Luckhart, S., 2008. Metabolic pathways in Anopheles stephensi mitochondria. Biochem. J 415, 309–316. 10.1042/BJ20080973

Goldstrohm, D.A., Pennington, J.E., Wells, M.A., 2003. The role of hemolymph proline as a nitrogen sink during blood meal digestion by the mosquito *Aedes aegypti*. J. Insect Physiol. 49, 115–121. 10.1016/s0022-1910(02)00267-6

Goncalves, R., Machado, A.C.L., Paiva-silva, G.O., Marcos, H.F., Momoli, M.M., Oliveira, J.H.M., Vannier-santos, M.A., Oliveira, P.L., Oliveira, M.F., 2009. Blood-Feeding Induces Reversible Functional Changes in Flight Muscle Mitochondria of *Aedes aegypti* Mosquito. PLoS One 4, 7854–7865. 10.1371/journal.pone.0007854

Hartwig, S., Kotzka, J., Lehr, S., 2015. Isolation and quality control of functional mitochondria. Methods Mol. Biol. 1264, 9–23. 10.1007/978-1-4939-2257-4_2

Hu, J., Liang, P., Shi, X., Gao, X., 2008. Effects of insecticides on the fluidity of mitochondrial membranes of the diamondback moth, Plutella xylostella, resistant and susceptible to avermectin. J. Insect Sci. 8, 3. 10.1673/031.008.0301

Jacobs, H.T., George, J., Kemppainen, E., 2020. Regulation of growth in Drosophila melanogaster: the roles of mitochondrial metabolism. J. Biochem. 167, 267–277. 10.1093/jb/mvaa002

Mailloux, R.J., 2015. Teaching the fundamentals of electron transfer reactions in mitochondria and the production and detection of reactive oxygen species. Redox Biology 4, 381–398. 10.1016/j.redox.2015.02.001

Mason, T. L., & Schatz, G. 1973. Cytochrome c oxidase from Bakers’ Yeast. In Journal of Biological Chemistry. Vol. 248, Issue 4, pp. 1355–1360. 10.1016/s0021-9258(19)44306-8

Merkey, A.B., Wong, C.K., Hoshizaki, D.K., Gibbs, A.G., 2011. Energetics of metamorphosis in Drosophila melanogaster. J. Insect Physiol. 57, 1437–1445. 10.1016/j.jinsphys.2011.07.013

Moosavi, B., Berry, E.A., Zhu, X.-L., Yang, W.-C., Yang, G.-F., 2019. The assembly of succinate dehydrogenase: a key enzyme in bioenergetics. Cell. Mol. Life Sci. 76, 4023–4042. 10.1007/s00018-019-03200-7

Nicholls, D.G., Palmeira, C.M., Moreno, A.J., 2012. Mitochondrial Bioenergetics: Methods and Protocols.

Pekny, J.E., Smith, P.B., Marden, J.H., 2018. Enzyme polymorphism, oxygen and injury: a lipidomic analysis of flight-induced oxidative damage in a succinate dehydrogenase d (Sdhd)-polymorphic insect. J. Exp. Biol. 221. 10.1242/jeb.171009

Palmer, C. S., Osellame, L. D., Stojanovski, D., & Ryan, M. T. 2011. The regulation of mitochondrial morphology: intricate mechanisms and dynamic machinery. Cellular Signalling, 23(10), 1534–1545. 10.1016/j.cellsig.2011.05.021

Palmeira, C. M., Rolo, A. P. 2012. Mitochondrial membrane potential (ΔΨ) fluctuations associated with the metabolic states of mitochondria. Mitochondrial Bioenergetics: Methods and Protocols, 89-101.

Picard, M., McEwen, B.S., 2018. Psychological Stress and Mitochondria. Psychosom. Med. 80, 141–153. 10.1097/PSY.0000000000000545

Popkov, V. A., Plotnikov, E. Y., Zorova, L. D., Pevzner, I. B., Silachev, D. N., Babenko, V. A., Jankauskas, S. S., Zorov, S. D., & Zorov, D. B. 2016. Quantification of mitochondrial morphology *in situ*. Tsitologiia, 58(9), 699–706. https://www.ncbi.nlm.nih.gov/pubmed/30198684

Pullman, M. E., Penefsky, H. S., Datta, A., & Racker, E. 1960. Partial resolution of the enzymes catalyzing oxidative phosphorylation. In Journal of Biological Chemistry (Vol. 235, Issue 11, pp. 3322–3329). 10.1016/s0021-9258(20)81361-1

Redecker, B., Zebe, E., 1988. Anaerobic metabolism in aquatic insect larvae: studies onChironomus thummi andCulex pipiens. J. Comp. Physiol. B 158, 307–315. 10.1007/BF00695329

Rodriguez-Morales, A., 2015a. Aedes: un eficiente vector de viejos y nuevos arbovirus (dengue, chikungunya y zika) en las Americas. Rev. cuerpo med. 8, 50–52.

Rodriguez-Morales, A., 2015b. No era suficiente con dengue y chikungunya: llegó también Zika. iMedPub Journals 11, 1–4. 10.3823/1245

Rodríguez, R., 2002. Estrategias para el control del dengue y del *Aedes aegypti* en las Américas. Rev. Cubana Med. Trop. 54, 189–201.

Rozeboom, L.E., 1960. *Aedes aegypti* (L.). The Yellow Fever Mosquito. Its Life History, Bionomics and Structure.Rickard Christophers. The Quarterly Review of Biology. 10.1086/403134

Saks, V.A., Belikova, Y.O., Kuznetsov, A.V., 1991. In vivo regulation of mitochondrial respiration in cardiomyocytes: specific restrictions for intracellular diffusion of ADP. Biochim. Biophys. Acta 1074, 302–311. 10.1016/0304-4165(91)90168-g

Saks, V.A., Veksler, V.I., Kuznetsov, A.V., Kay, L., Sikk, P., Tiivel, T., Tranqui, L., Olivares, J., Winkler, K., Wiedemann, F., Kunz, W.S., 1998. Permeabilized cell and skinned fiber techniques in studies of mitochondrial function in vivo. Mol. Cell. Biochem. 184, 81–100.

Salin, K., Villasevil, E.M., Anderson, G.J., Selman, C., Chinopoulos, C., Metcalfe, N.B., 2018. The RCR and ATP/O Indices Can Give Contradictory Messages about Mitochondrial Efficiency. Integr. Comp. Biol. 58, 486–494. 10.1093/icb/icy085

Singer, T.P., 1974. Determination of the activity of Succinate, NADH, Choline and a-Glycerophosphate dehydrogenases. Methods Biochem. Anal. 22, 123–175.

Slocinska, M., Barylski, J., Jarmuszkiewicz, W., 2016. Uncoupling proteins of invertebrates: A review. IUBMB Life 68, 691–699. 10.1002/iub.1535

Somlo, M., 1965. Induction des lacticocytochrome c reductases (D-ET L-) de la levure aerobie par les lactates (D-ET t-). Biochimica et Biophysica Acta - Bioenergetics 4, 183–201.

Thompson, S. N. (2003). Trehalose—the insect “blood”sugar. Advances in Insect Physiology. https://books.google.com/books

VenkatRao, V., Chaitanya, R.K., Naresh Kumar, D., Bramhaiah, M., Dutta-Gupta, A., 2016. Developmental and hormone-induced changes of mitochondrial electron transport chain enzyme activities during the last instar larval development of maize stem borer, Chilo partellus (Lepidoptera: Crambidae). Gen. Comp. Endocrinol. 239, 32–39. 10.1016/j.ygcen.2015.12.015

Vercesi, A.E., Bernardes, C.F., Hoffmann, M.E., Gadelha, F.R., Docampo, R., 1991. Digitonin permeabilization does not affect mitochondrial function and allows the determination of the mitochondrial membrane potential of Trypanosoma cruzi in situ. Journal of Biological Chemistry. 10.1016/s0021-9258(18)98703-x

Vital, W.D.E.O., 2006. Metabolismo de Glicose Durante a Embriogênese do Mosquito *Aedes aegypti* (Diptera: Culicidae).

Weaver, S., Lecuit, M., 2015. Chikungunya Virus and the Global Spread of a Mosquito-Borne Disease. The New England Jounal of Medicinenal of Medicine 372, 1231–1239. 10.1056/NEJMra1406035

Weinbach, E.C., Elwood Claggett, C., 1961. Oxidative Phosphorylation with Endogenous Substrates of Mitochondria. Journal of Biological Chemistry. 10.1016/s0021-9258(18)64208-5.

