## Supplementary material for "Baseline bioenergetic profile of mitochondria and permeabilized tissue of *Aedes aegypti* (Diptera: Culicidae) throughout its life cycle": Suplementary material.docx

**Figure 1.**

Average oxygen consumption (pmol/(sec*mL)) at stage 3 of mitochondria isolated from larvae, pupae, and thorax of *Aedes aegypti* females for the oxidizable substrates glutamate, malate, proline+pyruvate, succinate, and G3P. * Significant statistical differences between larvae, pupae, and adults for each oxidizable substrate evaluated: glutamate; malate; proline+pyruvate; succinate, and G3P (ANOVA test p≤0.05).

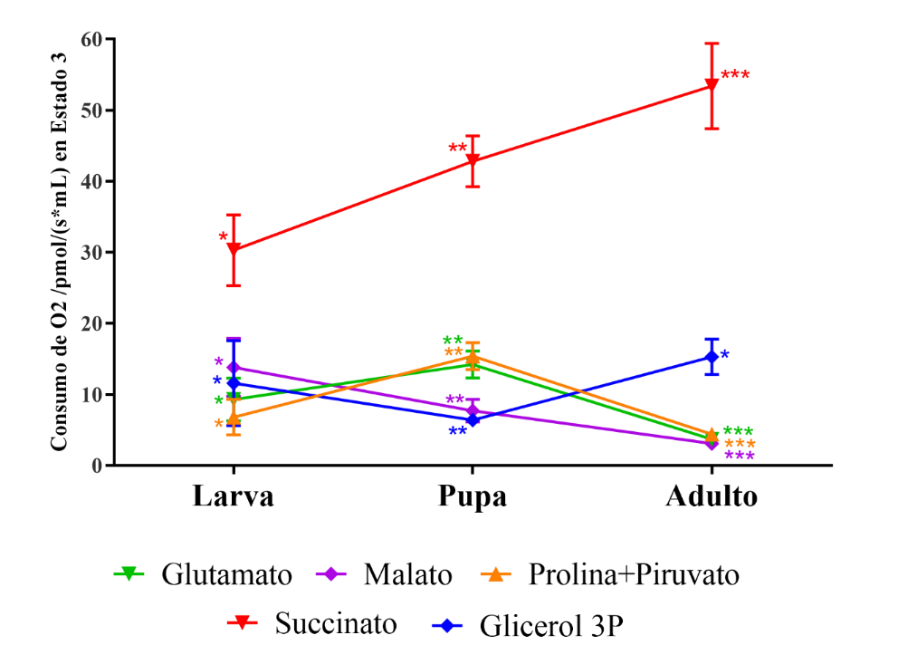

**Table 1.**

Oxygen consumption records (pmol/(sec*mL)) of isolated mitochondria and permeabilized individuals for L3-4 larvae, pupae, and thorax of *Aedes aegypti* females for the oxidizable substrates glutamate, malate, proline+pyruvate, succinate, and G3P.

Attached file in Excel format.
